# Does deterministic coexistence theory matter in a finite world?

**DOI:** 10.1101/290882

**Authors:** Sebastian J. Schreiber, Jonathan M. Levine, Oscar Godoy, Nathan J.B. Kraft, Simon P. Hart

## Abstract

Contemporary studies of species coexistence are underpinned by deterministic models that assume that competing species have continuous (i.e. non-integer) densities, live in infinitely large landscapes, and coexist over infinite time horizons. By contrast, in nature species are composed of discrete individuals subject to demographic stochasticity, and occur in habitats of finite size where extinctions occur in finite time. One consequence of these discrepancies is that metrics of species coexistence derived from deterministic theory may be unreliable predictors of the duration of species coexistence in nature. These coexistence metrics include invasion growth rates and niche and fitness differences, which are now commonly applied in theoretical and empirical studies of species coexistence. Here we test the efficacy of deterministic coexistence metrics on the duration of species coexistence in a finite world. We introduce new theoretical and computational methods to estimate coexistence times in stochastic counterparts of classic deterministic models of competition. Importantly, we parameterized this model using experimental field data for 90 pairwise combinations of 18 species of annual plants, allowing us to derive biologically-informed estimates of coexistence times for a natural system. Strikingly, we find that for species expected to deterministically coexist, habitat sizes containing only tens of individuals have predicted coexistence times of greater than 1, 000 years. We also find that invasion growth rates explain 60% of the variation in intrinsic coexistence times, reinforcing their general usefulness in studies of coexistence. However, only by integrating information on both invasion growth rates and species’ equilibrium population sizes could most (> 99%) of the variation in species coexistence times be explained. This integration is achieved with demographically uncoupled single species models solely determined by the invasion growth rates and equilibrium population sizes. Moreover, because of a complex relationship between niche overlap/fitness differences and equilibrium population sizes, increasing niche overlap and increasing fitness differences did not always result in decreasing coexistence times as deterministic theory would predict. Nevertheless, our results tend to support the informed use of deterministic theory for understanding the duration of species coexistence, while highlighting the need to incorporate information on species’ equilibrium population sizes in addition to invasion growth rates.

## Introduction

Understanding how competing species coexist is a central problem in ecology [Hutchinson, 1961, Chesson, 2000a]. Recent theoretical progress on this problem has replaced vague conceptualizations of coexistence requirements with tools that allow ecologists to quantify the niche differences that allow coexistence, and the fitness differences that drive competitive exclusion [Chesson, 1990, 2000a, Adler et al., 2007]. This progress has, in turn, motivated a large number of empirical studies to apply tools derived from theory to quantify the drivers of species coexistence in the field [e.g., Levine and Hille Ris Lambers, 2009, Narwani et al., 2013]. Importantly, however, the deterministic theory on which these advances are based assumes that competition occurs between populations whose densities vary continuously on landscapes of infinite size [Faure and Schreiber, 2014, Schreiber, 2017]. Under these assumptions, the influence of processes occurring as a consequence of the discrete nature of individuals–such as demographic stochasticity–are excluded [Hart et al., 2016, Pande et al., 2020b]. This generates a fundamental disconnect between theory and reality because while theory predicts that coexisting species will coexist indefinitely, in nature coexistence can only occur over finite periods of time. How well metrics derived from deterministic coexistence theory predict the duration of coexistence in the discrete, finite systems of nature – the ultimate object of study – remains largely unknown.

One of the most widely used metrics in contemporary theoretical and empirical studies of species coexistence are invasion growth rates [Hofbauer and Sigmund, 1998, Chesson, 2000a, Schreiber, 2000, Grainger et al., 2019]. A metric from deterministic models, the invasion growth rate of a species is its per-capita growth rate at vanishingly low densities when its competitors’ densities are at equilibrium. For two competing species, deterministic models predict that coexistence occurs when each species in a competing pair has a positive long-term invasion growth rate [Macarthur and Levins, 1967]. Provided there are no Allee effects or positive frequency-dependence at low densities [Schreiber et al., 2019], meeting this “mutual-invasibility” criterion implies that coexistence occurs indefinitely. Invasion analyses have been particularly powerful in studies of species coexistence because model-specific expressions for the invasion growth rate can be used to derive expressions that quantify the magnitude of niche and fitness differences between species [Chesson, 1990, 2013, Hart et al., 2018]. Thus, the mutual invasibility criterion has become central to much of our current understanding of species coexistence. However, when finite populations with positive deterministic invasion growth rates are depressed to low numbers of discrete individuals, they may still fail to persist because the negative effects of demographic stochasticity lead to extinction [Hart et al., 2018, Pande et al., 2020b]. More generally, even when species have higher densities away from the invasion boundary, demographic stochasticity operating on competing populations of finite size ensures extinction in finite time [Reuter, 1961, Lande et al., 2003, Jagers, 2010, Schreiber, 2017].

Despite the dominance of coexistence theory that imposes continuously varying population densities and infinitely large landscapes on a discrete, finite world, the effect of demographic stochasticity on coexistence is receiving greater attention [Vellend, 2010, Shoemaker et al., 2020]. Perhaps the most prominent example is neutral theory, in which demographic stochasticity is the sole driver of competitive dynamics between species [Hubbell and Foster, 1986, Hubbell, 2001]. Like fixation times for neutral alleles [Ewens, 2012], neutral theory predicts that competing species can coexist for long periods of time when the community size is large relative to the number of species [Hubbell, 2001]. The well-known problem with neutral theory, however, is that in emphasizing the role of demographic stochasticity, the theory simultaneously excludes all deterministic processes governing species interactions [Vellend, 2010]. This exclusion includes species-level fitness differences that drive competitive exclusion, and niche differences that promote coexistence [Adler et al., 2007]. Thus, depending on the relative strength of these deterministic processes, coexistence times predicted by neutral theory (i.e. by the action of demographic stochasticity alone), will be either significant over- or under-estimates.

More recently, population theoretic studies that incorporate demographic stochasticity into traditionally deterministic models of competition have begun to emerge [Adler and Drake, 2008, Gómez-Corral and López García, 2012, Gabel et al., 2013, Kramer and Drake, 2014]. These studies first make the important point that interspecific competitive interactions cause extinction dynamics to be different to that predicted by single-species models, and they also demonstrate that stochasticity causes the identity of the winner in competition to be less-than-perfectly predicted by deterministic competition-model parameters [Gómez-Corral and López García, 2012, Gabel et al., 2013, Kramer and Drake, 2014]. In addition, and together with models of extinction processes applied to single-species dynamics, these studies also highlight the likely importance of variables not traditionally considered in contemporary studies of species coexistence [Gómez-Corral and López García, 2012, Kramer and Drake, 2014]. For example, in single species models, time to extinction depends critically on equilibrium population size[Boyce, 1992, Grimm and Wissel, 2004, Ovaskainen and Meerson, 2010]. If equilibrium population size is similarly important for the duration of species coexistence, then the sole focus on invasion growth rates as the primary arbiter of coexistence may be problematic.

Despite recent progress in this area, existing studies of the effects of demographic stochasticity on the duration of coexistence concentrate only on cases where competitive exclusion is deterministically ensured [Gómez-Corral and López García, 2012, Kramer and Drake, 2014]. What is missing, therefore, are assessments of the duration of coexistence when coexistence rather than exclusion is deterministically ensured, and how these durations relate to existing deterministic coexistence metrics (e.g. invasion growth rates, niche and fitness differences). The latter point is particularly important as long as the field continues to rely on deterministic theory to identify coexistence mechanisms, and to interpret empirical results [Ellner et al., 2019]. Indeed, Pande et al. [2020b] identify inadequacies in invasion growth rates derived from deterministic theory for predicting coexistence times of finite populations in fluctuating environments. They show that the environmentally-dependent distribution of species’ growth rates in fluctuating environments – not just the average ‘invasion growth rate’ - influences coexistence times. While this finding is important, it leaves the independent effects of environmental and demographic stochasticity on coexistence times unresolved. For example, for coexistence mechanisms not reliant on environmental fluctuations there is no distribution of environmentally-dependent population growth rates, and so it remains unclear how demographic stochasticity alone influences coexistence times. Moreover, there is considerable empirical attention on, and support for, the contribution to coexistence of fluctuation-independent coexistence mechanisms [Adler et al., 2010, Godoy and Levine, 2014, Chu and Adler, 2015, Ellner et al., 2016, Mordecai et al., 2016, Letten et al., 2017, Li et al., 2019, Muller-Landau and Visser, 2019, Wainwright et al., 2019, Zepeda and Martorell, 2019, Spaak and De Laender, 2020]. For example, Zepeda and Martorell [2019] found that coexistence of 17 grassland species is mostly due to large fluctuation-independent coexistence mechanisms. Hence, it is particularly important to understand the applicability of these fluctuation-independent mechanisms for communities influenced by demographic stochasticity in nature.

In this paper, we explore the duration of species coexistence in discrete, finite natural systems experiencing demographic stochasticity. In particular, we determine the relationship between the duration of coexistence in the presence of demographic stochasticity and commonly used metrics of coexistence derived from deterministic theory. To determine these relationships, we introduce the concept of the intrinsic coexistence time, a multispecies analogue of Grimm and Wissel [2004]’s intrinsic extinction time for single-species stochastic population models. This metric corresponds to the mean time to losing one or more of these species after the species were coexisting sufficiently long to exhibit relatively stationary population dynamics i.e. quasi-stationarity. Our assessment of this metric is based on novel mathematical and computational methods that allow us to derive explicit relationships between the quasi-stationary behavior of a stochastic model and the dynamics of its deterministic model counterpart [Faure and Schreiber, 2014]. Importantly, we ground our analytical approach using estimates of competition model parameters from 90 pairs of annual plant species competing on serpentine soils in the field [Godoy et al., 2014].

## Models and Methods

### Models and Methods Overview

Our models and methods are composed of two broad parts. First, we describe the model and its empirical parameterization, we introduce metrics of deterministic coexistence, and we describe new analytical methods for calculating coexistence times in the presence of demographic stochasticity. Second, we describe our methods for answering a series of questions about the relationship between deterministic coexistence and stochastic coexistence times using the empirically-parameterized model.

### Model

We base our analysis on an annual plant competition model [Beverton and Holt, 1957, Leslie and Gower, 1958, Watkinson, 1980], which is well studied analytically [Cushing et al., 2004], and does a good job of describing competitive population dynamics in plant communities in the field [Godoy and Levine, 2014]. In the deterministic model with no demographic stochasticity, the dynamics of species *i* (*i* = 1 or 2) can be expressed in terms of its density *n_i,t_* and its competitor’s density *n_j,t_* at time *t*:

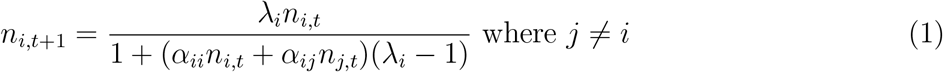

where *λ_i_* describes offspring production in the absence of competition, and *α_ii_* and *α_ij_* are the competition coefficients, which describe the rate of decline in offspring production as conspecific and heterospecific competitor densities increase, respectively. Including the multiplicative factor *λ_i_* — 1 in the denominator ensures that the competition coefficients are in units of proportional reductions in *λ_i_*, such that when *α_ii_n_i_* + *α_ij_n_j_* = 1 the population stops growing. Parameterizing our model in this way ensures that the conditions for coexistence in our model correspond to the classical conditions for coexistence in the continuous-time Lotka-Volterra competition model. In the deterministic model (1), offspring production and population densities take on values in the non-negative, real numbers.

The stochastic model takes the same functional form, but in contrast to the deterministic model, offspring production and population sizes *N_i,t_* take on non-negative integer values. Population densities *N_i,t_*/*S* are determined by the community size *S* and take on non-negative rational values. Specifically, in the stochastic model discrete individuals produce random numbers of discrete offspring according to a Poisson distribution. These random individual-level reproductive events generate demographic stochasticity in our model. Assuming the Poisson-distributed offspring production of individuals are independent with the same mean as the deterministic model, the sum of these events is also Poisson distributed and the dynamics of our stochastic model at the population level can be expressed as:

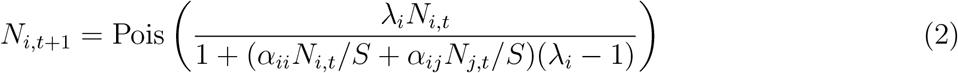

where Pois(*μ*) denotes a Poisson random variable with mean *μ*.

The community size parameter *S* allows us to quantify the effects of demographic stochasticity on coexistence in landscapes of finite size. Importantly, when community size *S* is sufficiently large, the dynamics of the densities *N_i,t_*/*S* of our stochastic model (2) are well approximated by the deterministic model (1) (Fig. 1A-C). This justifies our analysis of the effects of metrics of deterministic coexistence on stochastic coexistence times. Specifically, for any prescribed time interval, say [0, *T*], *N_i,t_*/*S* is highly likely to remain arbitrarily close to *n_i,t_* provided that *S* is sufficiently large and 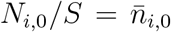 (Appendix S2). Intuitively, for fixed initial densities of both species, larger community sizes correspond to greater population sizes and, consequently, smaller stochastic fluctuations in their densities. However, in contrast to the deterministic model, both species eventually go extinct in finite time in the stochastic model (Appendix S2).

**Figure 1:**
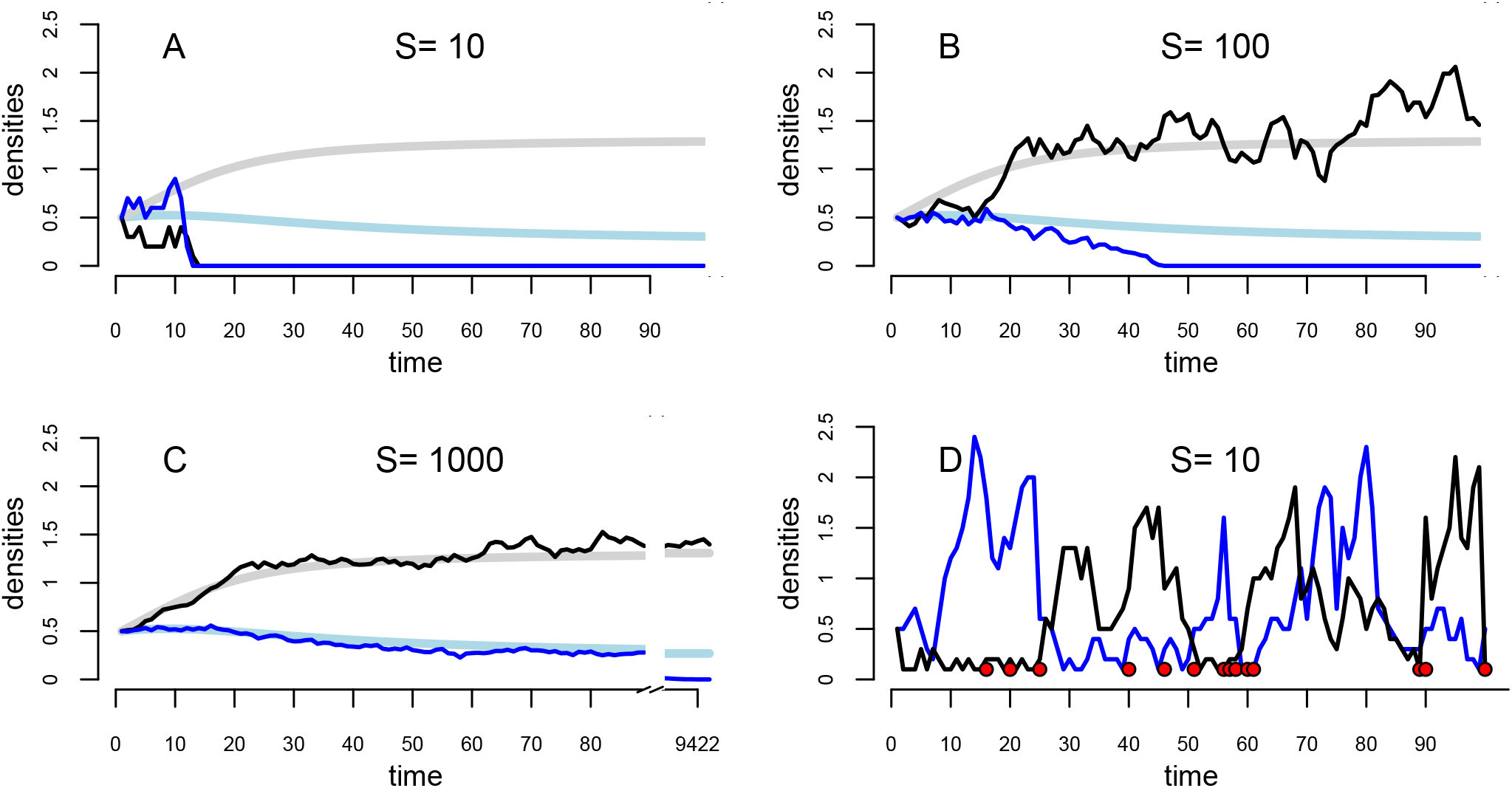
In panels A–C, the deterministic dynamics of the species densities *n_i,t_* for (1) are shown in light gray and light blue. Sample trajectories of the stochastic dynamics of the species densities *N_i,t_*/*S* for (2) are shown in blue and black. As the habitat size *S* increases, there is a closer correspondence between these dynamics, but eventually species go extinct in the stochastic model. In panel D, a sample simulation of the Aldous et al. [1988] algorithm to estimate the quasi-stationary distribution and the intrinsic coexistence time. Red circle correspond to times at which one or both species are going extinct; states at these times are replaced by states randomly sampled from the past. The frequency *p* of these red dots, for sufficiently long runs, determines the intrinsic coexistence time 1/*p*.

### Model parameterization

In our analyses we use parameter values for our competition model that were estimated for 90 pairwise combinations of 18 annual plant species competing in experimental field plots on serpentine soils in California [Godoy et al., 2014]. We used established methods to estimate these parameters [Hart et al., 2018]. Briefly, prior to the growing season we sowed focal individuals of each species into a density gradient of competitors. By recording the fecundity of each focal individual near the end of the growing season within a competitive neighborhood, we were able to estimate per germinant seed production in the absence of competition, and the rate of decline in seed production as competitor density increases [Hart et al., 2018]. We also measured the germination rate of each species and seed survival rate in the seedbank. See Godoy et al. [2014] for full details of our empirical and statistical methods.

These experiments were used to parameterize a model of the form

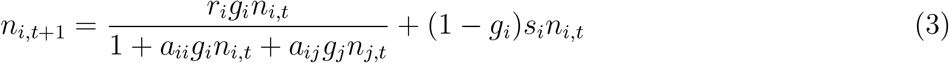

where *n*_*i,t*+1_ is the density of seeds in the soil after seed production but prior to germination, *r_i_* is the per germinant seed production in the absence of competition, *a_ii_* and *a_ij_* are the competition coefficients for germinated individuals, *g_i_* is the fraction of germinating seeds and *s_i_* is seed survival. As including the seed bank greatly complicates our analysis, yet ignoring it would unfairly bias competitive outcomes in the system, we assume that seeds that ultimately germinate do so in the year after they are produced. This implies that higher seed survival rates ultimately increase the average annual germination rate, 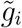 (Appendix S1). After making this adjustment, model (3) is equivalent to model (1) after setting *s* = 0, 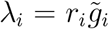 and 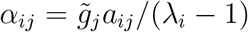. In the absence of a fluctuating environment (where seeds survival may buffer populations from extinction), this change is expected to have no effect on the duration of species coexistence

### Metrics of deterministic coexistence

Ultimately, we wish to relate coexistence times to metrics of deterministic coexistence commonly used in contemporary studies of species coexistence. Here, we focus on the invasion growth rate, and quantitative definitions of niche and fitness differences, which are themselves derivatives of the invasion growth rate. For (1), the invasion growth rate for species *i* is given by

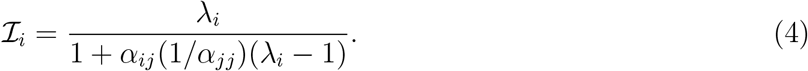

where 1/*α_jj_* is the single-species equilibrium density of species *j*. In the deterministic model, coexistence occurs if the invasion growth rates of both species are greater than one. Equivalently, this occurs when the minimum of the invasion growth rates, 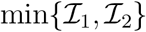, is greater than one. Conversely, if 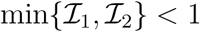, then one of the species’ densities will approach zero over the infinite time horizon.

Each species’ invasion growth rate is determined by: 1) frequency-dependent processes that provide growth-rate advantages to both species when they are at low relative density, and 2) frequency-independent processes that always favor the growth of one species over another regardless of relative density. These contributions to invasion growth rates have been quantitatively formalized as the niche overlap (*ρ*) and the average fitness ratio 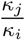 of the two species, respectively (see [Chesson, 2013] for mathematical details):

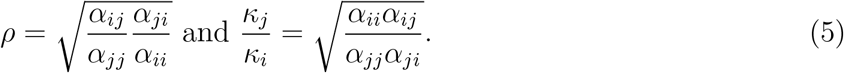

Niche overlap, *ρ*, decreases as the strength of intraspecific competition increases relative to the strength of interspecific competition. Low niche overlap causes species at high relative density to limit their own growth through intraspecific competition more than they limit the growth of species at low relative density through interspecific competition, leading to higher invasion growth rates. The fitness ratio 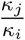 increases as one species becomes more sensitive to the total amount of competition in the system. For example, when 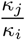 is greater than one, species *i* is more impacted by competition and will be displaced by species *j* when there is perfect niche overlap. The fitness ratio is used to quantify fitness differences: the greater the magnitude of the log fitness ratio, the greater the fitness difference. Ultimately, both species have positive invasion growth rates when *ρ* < *κ_j_*/*κ_i_* and *ρ* < *κ_i_*/*κ_j_*, demonstrating that deterministic coexistence occurs when niche overlap is less than 1 and small relative to the fitness difference.

If the species deterministically coexist they will approach a globally stable equilibrium, 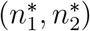. Solving for the densities at which both species’ fitness equals 1 in the deterministic model (1) gives

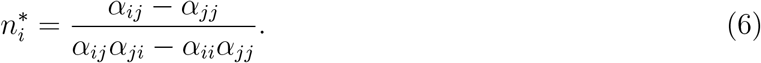

Invasion growth rates 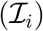 and equilibrium densities 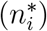 can also be expressed in terms of the niche overlap and the fitness ratio:

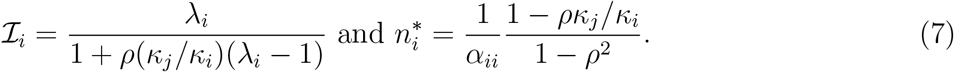

These expressions will become informative for interpreting the effects of niche overlap and fitness differences on coexistence times.

### Estimating intrinsic coexistence times

Unlike in the deterministic model, in the stochastic model both species always go extinct in finite time. We define the length of time prior to the first species going extinct as the coexistence time. The distribution of coexistence times in the stochastic model will depend on the initial conditions of the system. In studies of single-species persistence, Grimm and Wissel [2004] defined a quantity, the intrinsic mean time to extinction, that provides a common approach for selecting the initial distribution of the population and, thereby, allows for cross-parameter and cross-model comparisons. Here, we extend this work to introduce an analogous concept, the intrinsic coexistence time. The intrinsic coexistence time assumes that competing species have coexisted for a sufficiently long period of time in the past to exhibit relatively stationary population dynamics i.e. quasi-stationarity. At quasi-stationarity, the distribution of community population sizes is given by the model’s quasi-stationary distribution (QSD) [Méléard and Villemonais, 2012]. While in this quasi-stationary state, there is a constant probability, call it the persistence probability *p*, of losing one or both species each time step. We call the mean time 1/*p* to losing at least one of the species, the *intrinsic coexistence time*.

More precisely, the QSD for our stochastic model is a probability distribution *π*(*N_i_*,*N_j_*) on positive population sizes *N_i_* > 0, *N_j_* > 0 such that if the species initially follow this distribution (i.e. Pr[*N_i_*(0) = *N_i_*, *N_j_*(0) = *N_j_*] = *π*(*N_i_*, *N_j_*) for all *N_i_* > 0, *N_j_* > 0), then they remain in this distribution provided neither goes extinct i.e.

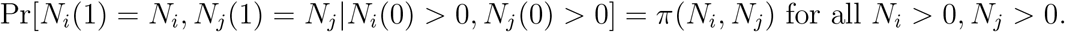

By the law of total probability, the persistence probability *p* when following the QSD satisfies

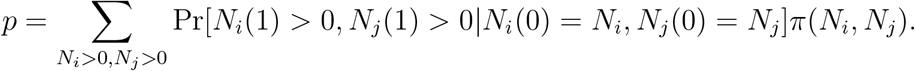

From a matrix point of view, *π* corresponds to the dominant left eigenvector of the transition matrix for the stochastic model and *p* is the corresponding eigenvalue [Méléard and Villemonais, 2012].

As the state space for the stochastic model is large even after truncating for rare events, directly solving for the dominant eigenvector is computationally intractable. Fortunately, there is an efficient simulation algorithm for approximating the QSD due to Aldous et al. [1988]. This algorithm corresponds to running a modified version of the stochastic model (Fig. 1D). Whenever one of the species goes extinct in the simulation, the algorithm replaces this state with a randomly sampled state from the past. More precisely, if (*Ñ_i_*(1), *Ñ_j_*(1)),…, (*Ñ_i_*(*t*), *Ñ_j_*(*t*)) are the states of the modified process until year *t*, then compute (*Ñ_i_*(*t* + 1), *Ñ_j_*(*t* + 1)) according to (2). If *Ñ_i_*(*t* + 1) = 0 or *N_j_*(*t* + 1) = 0, then replace (*Ñ_i_*(*t* + 1), *Ñ_j_*(*t* + 1)) by randomly choosing with equal probability 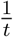 amongst the prior states (*Ñ_i_*(1), *Ñ_j_*(1)),…, (*Ñ_i_*(*t*), *Ñ_j_*(*t*)). The empirical distribution of (*Ñ_i_*(1), *Ñ_j_*(1)),…, (*Ñ_i_*(*t*), *Ñ_j_*(*t*)) approaches the QSD as *t* → ∞. This algorithm converges exponentially fast to the QSD [Benaïm and Cloez, 2015]. We use this algorithm to estimate coexistence times in our analyses. In Appendix S8, we also show how this algorithm also applies to a large class of models simultaneously accounting for demographic, environmental stochasticity, and spatial heterogeneity.

Having introduced our model and methods for determining coexistence times, we now describe how we apply these methods to address a series of questions about the relationship between deterministic coexistence and stochastic coexistence times for our empirically-parameterized models. We emphasize that while our models are empirically parameterized and so grounded in real biology, the focus of our analysis is a comparison between the predictions of the empirically-parameterized deterministic model and the predictions of the empirically-parameterized stochastic model. Understanding the relationship between our predicted coexistence times and observed coexistence times in nature is exceedingly difficult, likely requiring empirical studies over hundreds to thousands of generations in most systems. This is beyond the scope of our current analysis.

### How do deterministic coexistence and competitive exclusion relate to coexistence times?

We first explore how deterministic coexistence and deterministic competitive exclusion are related to coexistence times in the presence of demographic stochasticity. We do this through a mixture of analytical and numerical approaches. The analytical approach uses large deviation theory [Faure and Schreiber, 2014] to characterize how intrinsic coexistence times scale with community size *S* and how this scaling depends on whether the deterministic model predicts coexistence or exclusion. Based on our empirical parameterizations, competition between 82 species pairs is expected to result in deterministic competitive exclusion (i.e. parameter values result in 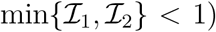, while 8 species pairs are expected to stably coexist (i.e. parameter values result in 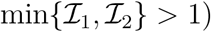. The remaining species pairs were excluded from our analyses either because estimates for at least one of the model parameters were missing or because one of the species had an intrinsic fitness *λ_i_* of less than one. For the two groups of species pairs we focus on (i.e. either deterministically coexisting or resulting in deterministic exclusion), we used simulations of ten million years to compute intrinsic coexistence times *C* across a range of community sizes *S*. We describe the relationship between community size and coexistence times for the models of the species pairs in each group.

### Do invasion growth rates predict coexistence times?

We used linear regression to determine if deterministic invasion growth rates (4) influence intrinsic coexistence times. For this analysis, we calculated intrinsic coexistence times only for the species pairs that are expected to deterministically coexist. To estimate intrinsic coexistence times for these species, we set the community size *S* = 0.04. This size would be empirically justified for the smallest serpentine hummocks in our study landscape (those with a few m^2^ of suitable habitat), which might contain as few as 20 individuals of the subdominant species (based on germinable densities projected from Gilbert and Levine [2013]). The species that compose the focal pairs in this analysis tend to be more common and are often found on larger hummocks, but this small habitat size allows us to evaluate the dynamics of systems where the effects of demographic stochasticity might be substantial. For each species pair we used simulations of 10^7^ years to estimate the coexistence times predicted by the models. For seven of the species pairs predicted to coexist by the deterministic models, we were able to generate good estimates of intrinsic coexistence times predicted by the stochastic models because there were multiple extinction events in simulations of this length. For one species pair predicted to coexist by the deterministic model, there were no extinction events in the numerical simulations. Because we only have a lower bound of > 10^7^ years for the mean intrinsic coexistence time for this pair, it was excluded from our analysis.

Coexistence time was the dependent variable in our regression, and the log of 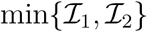 was the independent variable. We used this independent variable because the species with the lower invasion growth rate is, all else being equal, more likely to go extinct first. To examine the robustness of our conclusions, we repeated our analysis for 1, 000 randomly-drawn parameter values with community size *S* = 10. These random draws were performed on uniform distributions with [1.1, 2] for the *λ_i_* values, [0.2, 0.5] for the *α_ii_* values, and *α_ii_* × [0,1] for the *α_ji_* values (to ensure deterministic coexistence).

### Do equilibrium population sizes predict coexistence times?

Invasion growth rates were a less-than-perfect predictor of coexistence times for the stochastic models (see Results). Therefore, we also used linear regression to explore the relationship between the equilibrium population sizes of the coexisting species and the intrinsic coexistence time. For this analysis, we used the same methods as described above, but with the log of the minimum of the two equilibrium population sizes i.e. 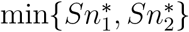 as the independent variable in the linear regression. To examine the robustness of our conclusions, we repeated our analysis for 1, 000 randomly-drawn parameter values with community size *S* = 10 as described in the section *Do invasion growth rates predict coexistence times?*, above.

### Do greater niche overlap or greater fitness differences always reduce coexistence times?

According to deterministic theory, greater niche overlap and greater fitness differences both have a negative impact on coexistence [May, 1975, Chesson, 2000a, Adler et al., 2007]. It is important, therefore, to understand if these negative effects on deterministic coexistence also have predictably negative effects on the duration of species coexistence. We investigate these relationships by calculating coexistence times as a function of niche overlap and fitness differences.

Our goal is to assess the independent effects of these determinants of coexistence on coexistence times. However, because niche overlap and fitness differences are both functions of the same parameters (5), they are not quantitatively (or biologically) inherently independent [see also Song et al., 2019]. This precludes using the natural variation in niche overlap and fitness differences observed across our focal species pairs for our analyses. Therefore, to achieve our goal, we manipulated niche overlap and fitness differences separately for each species pair. Specifically, to assess the effects of niche overlap on model predicted coexistence times, we multiplied the interspecific competition coefficients *α*_12_, *α*_21_ for each coexisting species pair by a common, fixed factor. This manipulation allows the niche overlap to vary while keeping the fitness ratio constant according to (5). Mechanistically, this may be interpreted as altering the degree to which the two species use the same resources. Similarly, to assess the effects of the fitness difference independent of any change in niche overlap, we multiplied the competition coefficients *α*_22_, *α*_21_ within each coexisting pair by a fixed factor. This manipulation reduces the sensitivity of species 2 to competition, increasing its competitive ability according to (5), while having no effect on niche overlap. Mechanistically, this may be interpreted as increasing the efficiency with which species 2 uses shared resources. Based on these manipulations, for each coexisting species pair we explored the relationship between niche overlap and fitness differences, and coexistence times. We interpret these relationships via the effects of niche overlap and fitness differences on invasion growth rates and equilibrium population sizes as per (7).

### Do stochastic competitive dynamics influence coexistence times?

Competition may also influence coexistence times because the stochastic population dynamics of the two species are coupled. For example, a stochastic increase in the population size of one species might be expected to result in a concomitant decrease in the population size of its competitor, increasing its risk of extinction. The effect on coexistence times of this dynamic coupling of the two species is not accounted for by static metrics of coexistence such as invasion growth rates and equilibrium population sizes. To identify whether this dynamic coupling plays an important role in determining coexistence times, we built a simplified model, what we call the demographically uncoupled model, incorporating the effects of competition on invasion growth rates and equilibrium population sizes, but excluding the coupling of the stochastic fluctuations in the population sizes of the competitors. In our demographically uncoupled model, each species has a low density growth rate “*λ_i_*” and carrying capacity “1/*α_ii_*” that matches its invasion growth rate 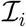 and equilibrium density 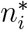, respectively, from the deterministic two-species model. The update rule for this simplified model is

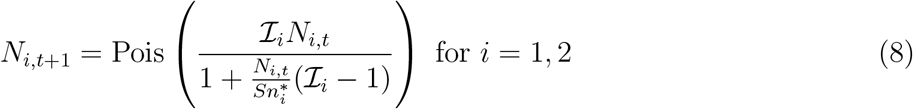

Importantly, note that the change in population size of *N_i_* does not depend on *N_j_*. Therefore, in the demographically uncoupled model the effects of interspecific competition on invasion growth rates and equilibrium population sizes are retained, but the species are dynamically uncoupled. By quantifying the relationship between coexistence times calculated for the full (2) and demographically uncoupled (8) models, we can determine whether the combined effects of competition on invasion growth rates and equilibrium population sizes are sufficient to predict coexistence times, or if competitive dynamics arising from the coupled stochastic dynamics of the species are also important.

In Appendix S7, we show the coexistence times for the demographically uncoupled model reduce to calculating the intrinsic persistence times *P_i_* of each species in this simplified model. These persistence times *P_i_* are calculated independently for each species using the [Aldous et al., 1988] algorithm. Then, the coexistence time *C*_simplifled_ of the demographically uncoupled model equals:

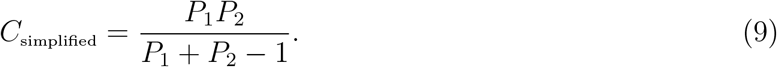

When the intrinsic persistence times *P_i_* for each species are sufficiently long (e.g. 10 years or more), the coexistence time predicted by this simplified model is approximately one half of the harmonic mean of the intrinsic persistence times for the two species: 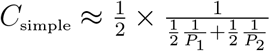. As the harmonic mean is dominated by the minimum of its arguments, the species with the shorter persistence time has the greater influence on the coexistence time.

We calculated intrinsic coexistence times for the full and demographically uncoupled models for the eight species pairs that are expected to deterministically coexist. We then used linear regression to determine the extent to which the simplified model explains coexistence times in the full model. To examine the robustness of our conclusions, we repeated our analysis for 1, 000 randomly-drawn parameter values with habitat size *S* = 10 as described in the section *Do invasion growth rates predict coexistence times?*, above.

## Results

### How do deterministic coexistence and competitive exclusion relate to coexistence times?

The duration of species coexistence tended to be several orders of magnitude larger for species expected to deterministically coexist than for those pairs where deterministic exclusion is expected (Fig. 2). Indeed, our numerical results demonstrate that species expected to deterministically coexist will tend to do so for decades to millennia, even when community sizes are small. By contrast, our numerical simulations suggest that the vast majority of the 82 species pairs for which deterministic exclusion is expected will tend to coexist for less than five years (Fig. 2A).

**Figure 2:**
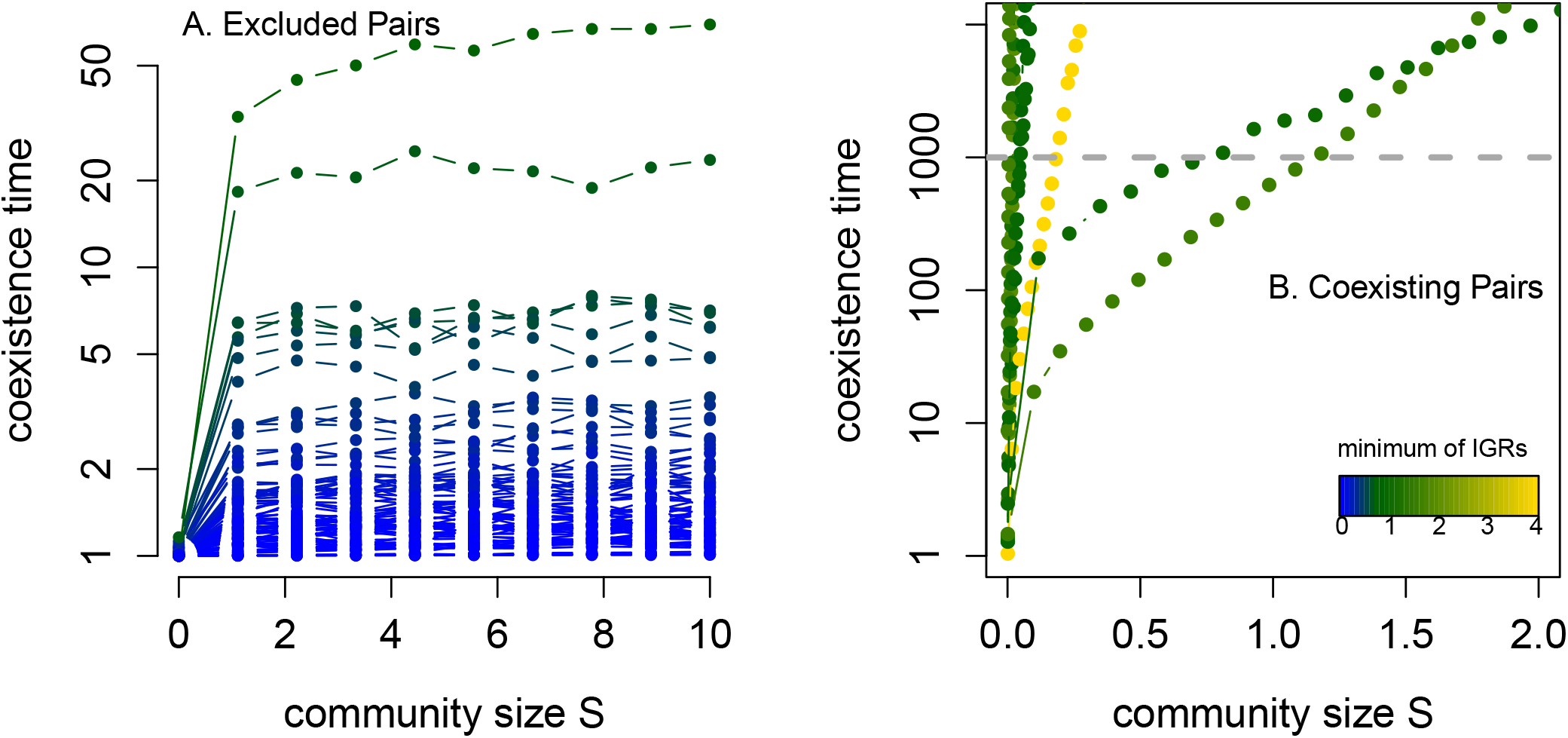
Coexistence times as a function of community size *S* for deterministically excluded species pairs (A) and deterministically coexisting pairs (B). For each viable species pair, the invasion growth rates 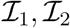 were computed for the associated deterministic model. For each species pair, the coexistence time of the associated individual-based model were calculated for a range of community sizes *S*. Each of these species pair curves are colored by their 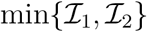 value.

Our analytical and numerical results expose a fundamental dichotomy between deterministic coexistence and exclusion in terms of how coexistence times for the models scale with increasing community size. In particular, if deterministic exclusion is predicted 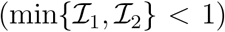, then our mathematical analysis (Appendix S3) implies that the intrinsic coexistence times are bounded above by 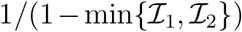 and do not increase significantly with community size. This analytical result is consistent with our numerical results for the 82 models associated with the species pairs for which deterministic exclusion was predicted (Fig. 2A). By contrast, if the two species are predicted to deterministically coexist 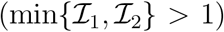, then our analysis implies that intrinsic coexistence times scale exponentially with the community size *S*. This means that small increases in community size lead to very large increases in the duration of species coexistence. This analytical result is also consistent with our simulations of the 8 species pairs for which deterministic exclusion was predicted (Fig. 2B).

For species pairs predicted to deterministically coexist, the mathematical analysis only ensures there is a constant **✶** > 0 such that the intrinsic coexistence time is proportional to *e*^**✶***S*^. There is no simple formula for **✶** as it will depend on many details of the individual-based model (see proofs in [Faure and Schreiber, 2014]). To understand some of these dependencies, we explored how coexistence times depend on key coexistence metrics from the deterministic model for a fixed community size *S* = 0.04, (a small hummock; see methods), the results of which we describe next.

### Do invasion growth rates predict coexistence times?

Across seven pairs of deterministically coexisting competitors, we found a positive correlation between the minimum invasion growth rate, 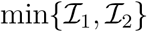, and intrinsic coexistence times, as determined from the quasi-stationary distributions of the empirically-parameterized model (adjusted *R*^2^ = 0.6038, *p* = 0.0244; Fig. 3A). Intuitively, the species with the lower invasion growth rate tends to go extinct first, and so it is the lower of the two invasion growth rates that predicts coexistence times. However, there was substantial variation in coexistence times unexplained by invasion growth rates, such that similar invasion growth rates resulted in several orders of magnitude difference in coexistence times (Fig. 3A). Random sampling of parameter space produced similar results (Appendix S4:Fig. S1A).

**Figure 3:**
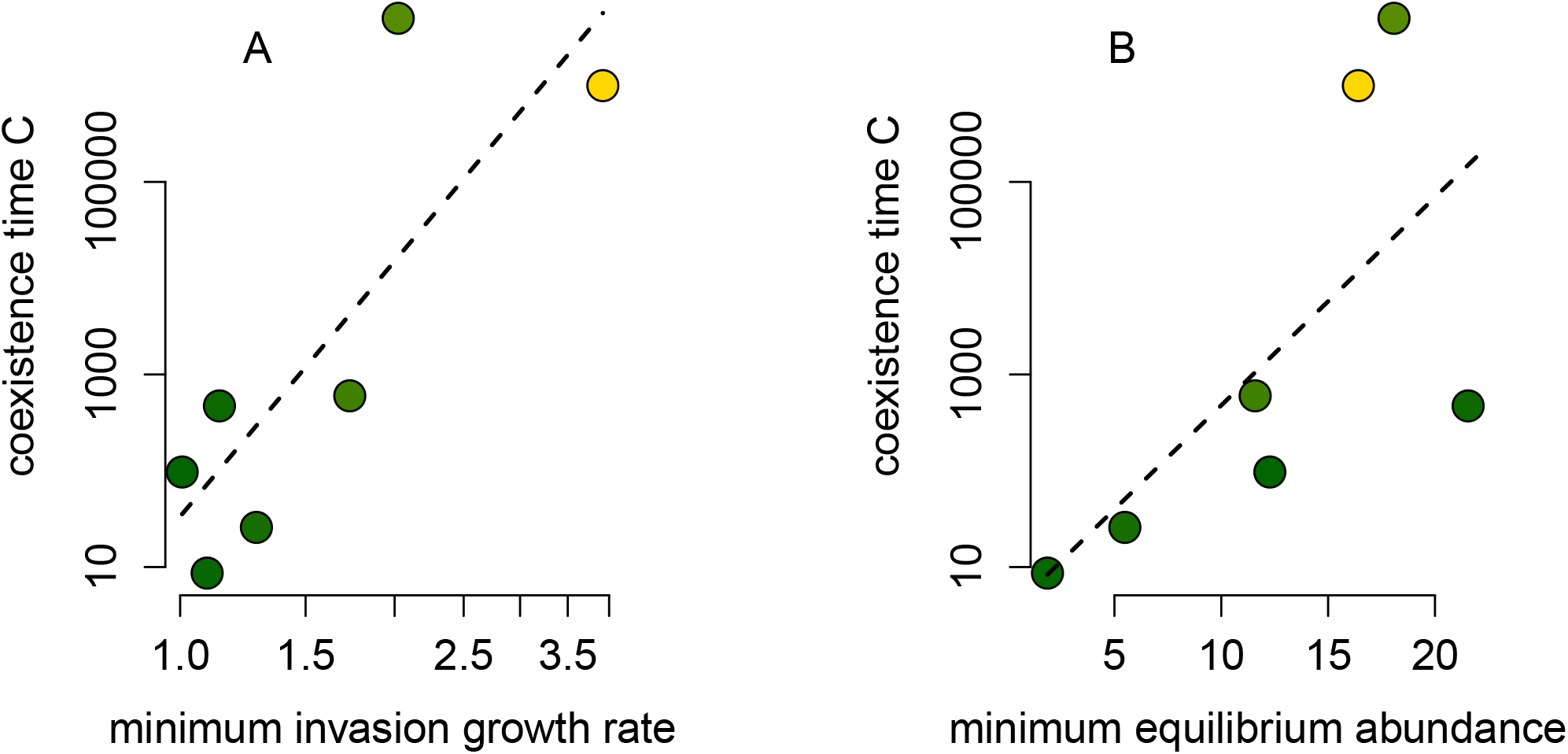
Predicting coexistence times using the minimum, 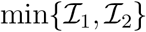, of invasion growth rates (A) and the minimum, 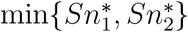, of equilibrium population sizes (B). Both panels plotted on a log–log scale. Each of these species pair curves are colored by their 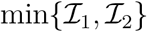 value as in Figure 2.

### Do equilibrium population sizes predict coexistence times?

We also found a positive correlation between the minimum of the equilibrium population sizes, 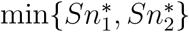, and coexistencetimes (adjusted *R*^2^ = 0.3386; *p* = 0.09964, Fig. 3B). As equilibrium population sizes did a worse job of explaining the variation in the coexistence times than the invasion growth rates, similar equilibrium population sizes also resulted in several orders of magnitude of difference in coexistence times. Random sampling of parameter space produced similar results (Appendix S4:Fig. S1B).

### Do greater niche overlap or greater fitness differences always reduce coexistence times?

Greater niche overlap and greater fitness differences did not always reduce coexistence times, as deterministic theory would predict (Figs. 4,5). For 5 out of 7 species pairs, both greater niche overlap and greater fitness differences reduced the coexistence times predicted by the models (Figs. 4A,D; Appendix S6). For these pairs, the relationships were nonlinear, but negatively monotonic. However, for the remaining two species pairs, the effects of greater niche overlap and fitness differences on coexistence times were complex (Figs. 5A,D; Appendix S6). For these species pairs, increasing niche overlap consistently reduced coexistence times, but in a highly nonlinear fashion (Fig. 5A). By contrast, increasing fitness differences had both positive and negative effects on coexistence times for the models associated with these species pairs, depending on the magnitude of the change in the fitness difference (Fig. 5D). Importantly, for the species pairs where these complex effects emerged, the competitively inferior species (as detemined by the fitness ratio in equation (5)), had the higher (i.e not the minimum) equilibrium population size.

**Figure 4:**
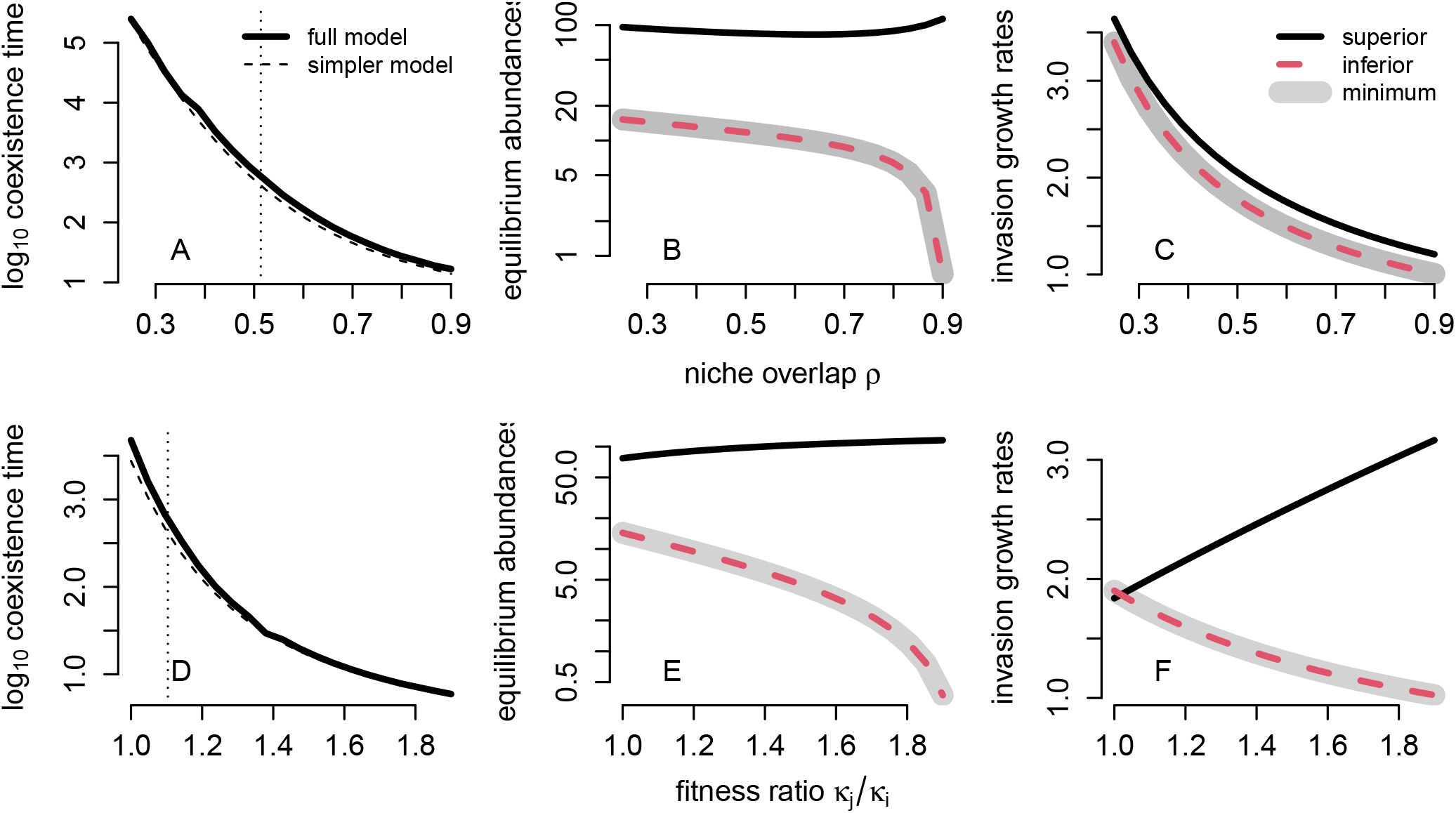
Effects of niche overlap and fitness ratios on coexistence times for a pair of competing species where the fitness inferior has the lower equilibrium abundance. In (A) and (B), log_10_ coexistence times and the demographically uncoupled model approximations (dashed lines) are plotted. The vertical dotted line corresponds to the base empirical value. In (B) and (E), the deterministic equilibrium population sizes 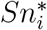 are plotted with the gray shading indicating the minimum of the two population sizes. In (C) and (F), the deterministic invasion growth rates of the deterministic, mean field model are plotted for the fitness inferior (dashed red) and fitness superior (solid black) with gray shading indicating the minimum of two invasion growth rates.

**Figure 5:**
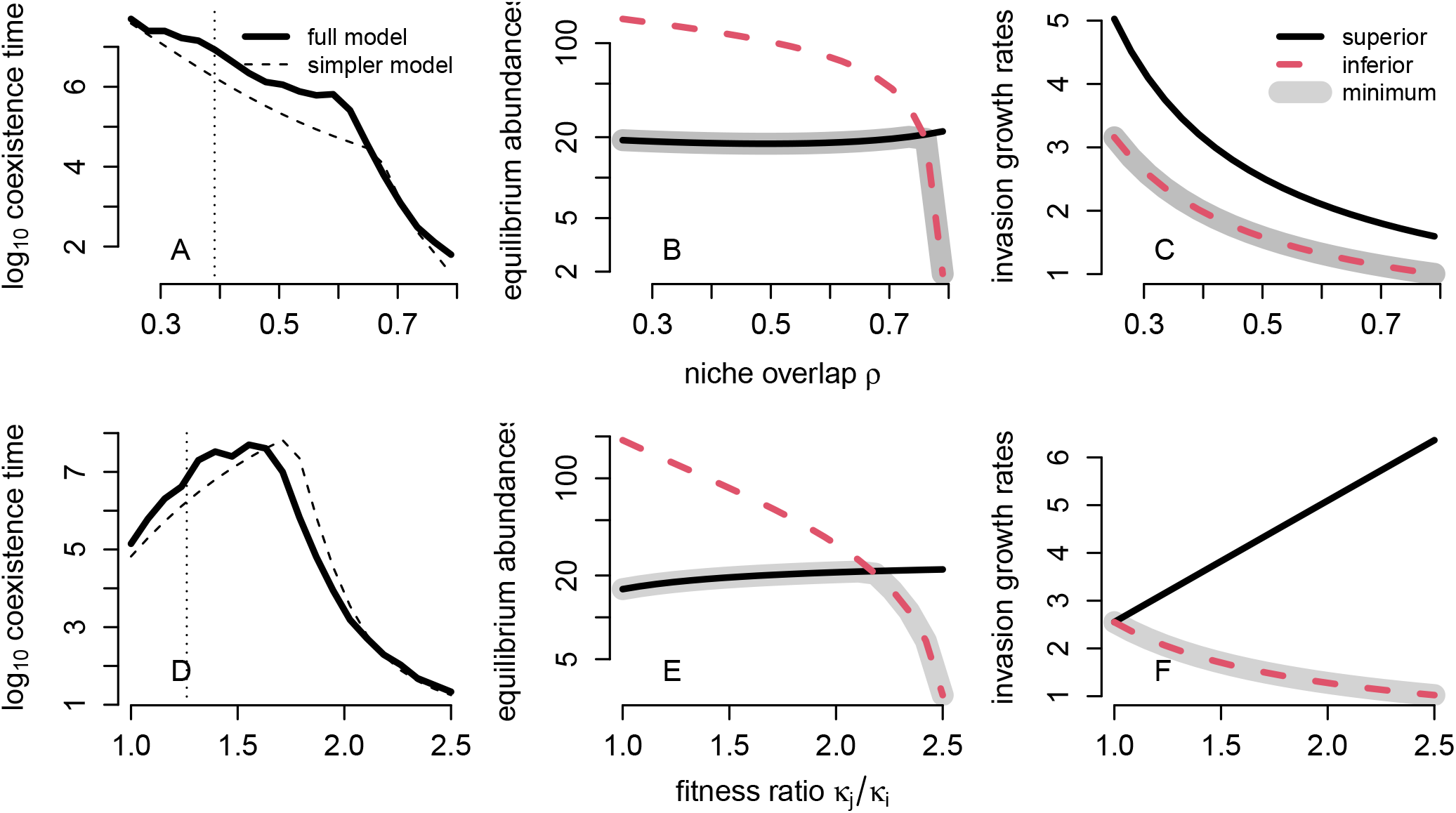
Effects of niche overlap and fitness ratios on coexistence times for a pair of competing species where the fitness superior has the lower equilibrium abundance. In (A) and (B), log_10_ coexistence times and the demographically uncoupled model approximations (dashed lines) are plotted. The vertical dotted line corresponds to the base empirical value. In (B) and (E), the deterministic equilibrium population sizes 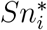 are plotted with the gray shading indicating the minimum of the two population sizes. In (C) and (F), the deterministic invasion growth rates of the deterministic, mean field model are plotted for the fitness inferior (dashed red) and fitness superior (solid black) with gray shading indicating the minimum of two invasion growth rates.

When the inferior competitor has the higher equilibrium population size, complex effects on coexistence times emerge via the influence of niche overlap and fitness differences on the minimum of the equilibrium population sizes and the minimum of the invasion growth rates (see (7)). For example, as fitness differences increase, the superior competitor’s equilibrium population size also increases (equation (7), Appendix S6, Fig. 5E,F). Consequently, when the superior competitor has the lower equilibrium population size, increasing the fitness difference increases the minimum of the equilibrium population sizes, which increases coexistence times (Fig. 5D). Increasing niche overlap can also increase the equilibrium population size of the superior competitor, but always decreases its invasion growth rate (equation (7), Appendix S6, Fig. 5B,C). When the superior competitor has the lower equilibrium abundance, these opposing trends result in coexistence times that decrease in a highly nonlinear manner with increasing niche overlap. A general result is that coexistence times decrease when both the minimum invasion growth rate and the minimum equilibrium population size decline with increasing niche overlap and increasing fitness differences (Figs. 4,5).

### Do stochastic competitive dynamics influence coexistence times?

To isolate the influence of coupled stochastic competitive dynamics on coexistence times we compared coexistence times calculated from the full model (2) including these coupled dynamics, to coexistence times calculated from the demographically uncoupled model (8) excluding these coupled dynamics. As described in the methods, the demographically uncoupled model is two uncoupled, individual-based single species models whose low-density growth rate and intraspecific competition coefficients equal 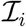 and 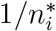, respectively. For the 7 species pairs predicted to coexist in the deterministic model, the simplified model incorporating competition only through its effects on invasion growth rates and equilibrium population sizes did an exceptional job in predicting the actual coexistence time (log *C* = 1.031 log *C*_simple_ with *R*^2^ = 0.9977 and *p* < 10^-8^, Fig. 6). Random sampling of parameter space produced similar results (Appendix S4:Fig. S1C).

**Figure 6:**
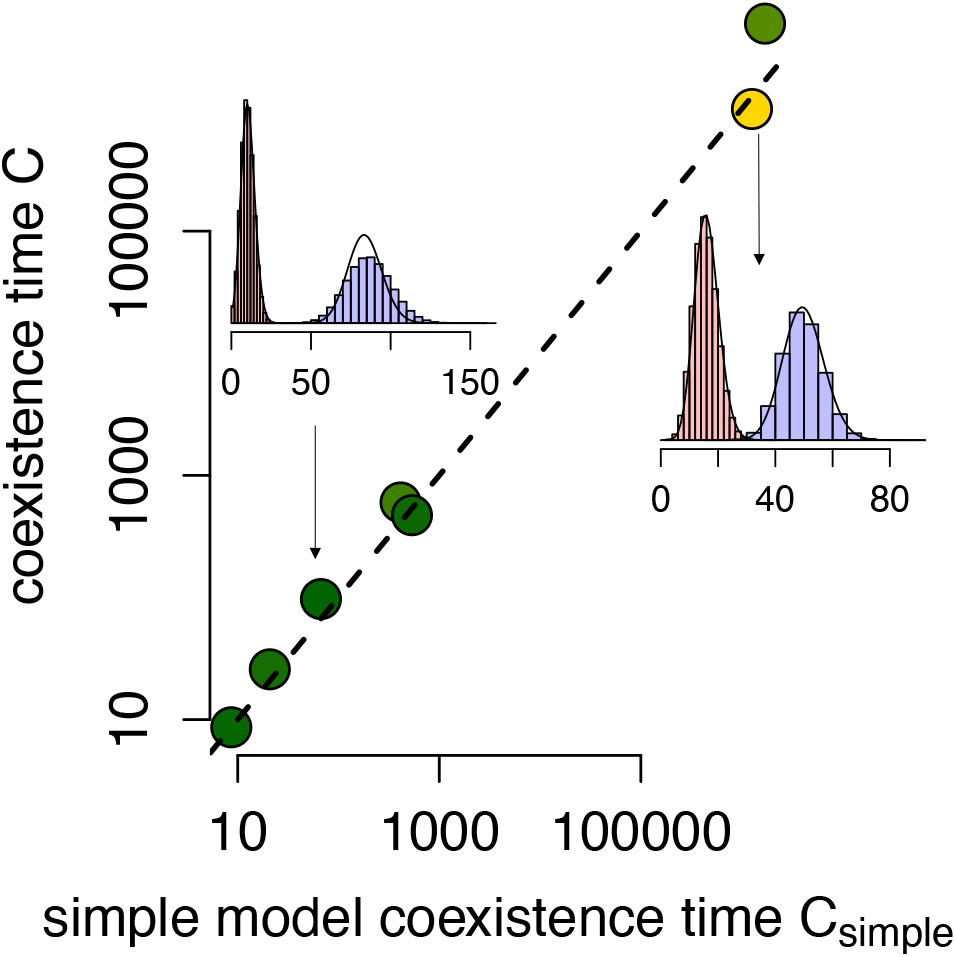
Predicting coexistence times with the demographically uncoupled model (8). Dashed line is the 1-to-1 line. The main panel shows how the demographically uncoupled models incorporating only invasion growth rates and equilibrium population sizes predict coexistence times for the full, demographically coupled models. Insets: Bar plots show the simulation-based estimates of the coupled, competitive model’s quasi-stationary distributions and the black curves are the analytical estimates from the demographically uncoupled model. Each of these species pair curves are colored by their 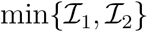 value as in Figure 2.

## Discussion

We introduced a new metric, the intrinsic coexistence time, that characterizes the risk of species loss due to demographic stochasticity over any time horizon. This metric complements more traditional coexistence metrics based on invasion growth rates and can be computed for many existing data-based models by following a two step procedure. First, extend the deterministic model to account for demographic stochasticity. Informing such a stochastic model with demographic data obtained in a field context is not much more difficult than parameterizing deterministic models. Aside from fecundity, most transition rates (i.e. survival, growth, dispersal probabilities) of the deterministic model directly transfer to the stochastic model. The fecundity distribution can be estimated from the raw data used to calculate mean fecundity in deterministic models, or one can assume that fecundity is Poisson distributed with the calculated mean. Second, estimate each intrinsic coexistence time by a single simulation using the algorithm of Aldous et al. [1988] (see Appendix S8 for extensions to a broad class of models). Using this two-step approach, we evaluated how well metrics from deterministic theory predict intrinsic coexistence times for 18 species of Californian annuals.

Consistent with the deterministic theory, we find invasion growth rates are, in general, good predictors of intrinsic coexistence times in models explicitly accounting for demographic stochasticity. Thus, reinforcing the general usefulness of these invasion growth rates in theoretical and empirical studies of species coexistence. Their usefulness stems from two conclusions of our study. First, when both species’ invasion growth rates are positive (i.e. deterministic coexistence is predicted), intrinsic coexistence times increase exponentially with community size. Thus, for a given minimum invasion growth rate, doubling community sizes quadruples coexistence times. Strikingly, for the 8 species pairs of serpentine annuals predicted to deterministically coexist, community sizes corresponding to only tens of individuals of the less common species were still sufficient to ensure predicted coexistence times of greater than 1, 000 years (Fig. 2) – a time frame well beyond most empirically-relevant scenarios and much greater than considered in most conservation studies [Meine, 1999]. Second, for a given community size, we found that the minimum of the invasion growth rates 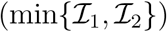 explained 60% of the variation in coexistence times of the serpentine annuals predicted to deterministically coexist (Fig. 3), and 82% of the variation of coexistence times for a random sampling of parameter space (Appendix S4:Fig. S1A). The two conclusions are consistent with recent work on coexistence times for the lottery model which simultaneously account for environmental stochasticity as well as demographic stochasticity [Pande et al., 2020b, Ellner et al., 2020]. The main differences being that invasion growth rates for the lottery model in the presence of temporal environmental stochasticity correspond to the geometric mean of fitness across time, with coexistence times increasing as a power law, rather than exponentially, with community size [Ellner et al., 2020].

Despite explaining a significant amount of variation, invasion growth rates are not perfect predictors of coexistence times in models accounting for demographic stochasticity. In particular, predicted coexistence times varied by an order of magnitude for species pairs of serpentine annuals with similar values of 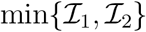 (Fig. 3). Thus, while higher invasion growth rates tend to be associated with longer coexistence times, and these coexistence times tend to be exceedingly long, the capacity of invasion growth rates to accurately predict coexist times should still be viewed with some caution. Pande et al. [2020b] raised a similar concern but for different reasons, when studying Chesson’s lottery model. They demonstrated that increases in environmental variability can simul-taneously lead to an increase in invasion growth rates and shorter coexistence times. Our study finds that some disassociation between invasion growth rates and coexistence times can occur even without temporal fluctuations in the fitness of the rare species.

Beyond invasion growth rates, we found that population sizes away from the invasion boundary (i.e. equilibrium population sizes) are also important determinants of coexistence times. This finding is consistent with single-species models of extinction risk, where both low-density growth rate and equilibrium population size determine risk of extinction [e.g., Lande et al., 2003]. However, in and of themselves, equilibrium population sizes were poorer predictors of coexistence times than invasion growth rates. The minimum of the equilibrium population sizes explained 26% less of variation than 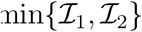 in the empirical species pairs (Fig. 3), and 12% less of the variation of the coexistence times in the random sampling of parameter space (Appendix S4:Fig. S1B).

When combined, the invasion growth rates and the equilibrium population sizes are such good determinants of coexistence times that simply knowing these two quantities can explain over 99% of the variation of the predicted coexistence times (Fig. 6, Appendix S4:Fig. S1C). Notably, this predictive capacity emerges from a demographically uncoupled model parameterized only with the invasion growth rates and equilibrium population sizes, while excluding the effects of the coupled stochastic dynamics of interspecific competition. The success of the demographically uncoupled model emphasizes that coexistence times in the presence of demographic stochasticity can be predicted using minimal information gleaned directly from an appropriate deterministic model. Whether this tractable approach applies to stochastic models with more species or different nonlinearities in the per-capita growth rates is an important direction for future research.

As the predicted coexistence time for the demographically uncoupled model is approximately the harmonic mean of the persistence time for each of the (re-calibrated) single-species models, one can use it to gain additional insights into coexistence times for the competing species. For example, if both competitors have large invasion growth rates (i.e. 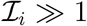), then each competitors’ persistence time is approximately equal to its exponentiated equilibrium population abundance, 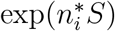 (see Appendix S7). In which case, for a given community size 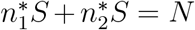, the predicted coexistence time is maximized when both species have equal equilibrium abundances (see Appendix S7). For a more diverse community, this is equivalent to when Shannon’s diversity index is maximized. However, if one species has a significantly lower invasion growth rate than its competitor, then maximizing the coexistence time requires that this species be over-represented in the community (i.e. the community would have a lower Shannon diversity index) (see Appendix S7).

Because invasion growth rates are imperfect predictors of coexistence times, coexistence metrics derived solely from invasion growth rates are also imperfect predictors of coexistence times. These coexistence metrics include modern quantitative definitions of niche overlap and fitness differences [Chesson, 2000a, 2013, Godoy and Levine, 2014, Chesson, 2018]. Because niche overlap and fitness differences can have opposing effects on the relative population sizes of the competitors, coexistence times do not necessarily decrease with increasing niche overlap or increasing fitness differences, as would be expected based only on deterministic theory (Fig. 5A,D). For models of two of the annual, serpentine species pairs, this unexpected outcome occurred when the inferior competitor had the higher equilibrium population size. Operationally, one can simply check whether this is the case before interpreting the effects of niche overlap and fitness differences as they are commonly derived and applied [Chesson, 2000, Adler et al., 2007, 2010, Gilbert and Levine, 2013, Chesson, 2013]. According to equation (7), the competitive inferior has the larger equilibrium abundance when it is insensitive to intraspecific competition (i.e. has a large carrying capacity 1/*α_jj_*) but highly sensitive to interspecific competition relative to the competitively superior species. This observation provides an indirect means of evaluating whether this inverted relationship is likely to occur in a given empirical system.

Our conclusions are based on an analysis of an annual plant competition model, which is a specific case of a large and more general class of models that can be re-expressed in Lotka-Volterra form. This class of models has been the focus of much of the theoretical and empirical work deriving and applying quantitative definitions of niche and fitness differences to studies of species coexistence [Macarthur and Levins, 1967, Chesson, 1990, 2000a, 2013, Godoy and Levine, 2014]. Given that niche and fitness differences in these models are derived from invasion criteria, we expect our conclusions about the effects of niche and fitness differences on coexistence times to apply across this class of models. More generally, equilibrium densities of competitors are likely to be important quantities in all models of competition regardless of their complexity, but are not often taken into account in recent assessments of coexistence. [Carroll et al., 2011, Chesson, 2013, Spaak and De Laender, 2020]. For example, [Spaak and De Laender, 2020] provided a general method for defining niche and fitness differences using per-capita growth rates evaluated at densities where at least one species is absent. However, these metrics do not take into account, and are unlikely to correlate with, coexistence equilibrium densities. Therefore, coexistence times are unlikely to map intuitively to niche and fitness differences even for these more general definitions. In sum, our work suggests that any coexistence metric not explicitly taking into account densities at which the species coexist deterministically may not predict coexistence times correctly.

For models considered here, all conspecific individuals have the same invasion growth rate across time – there is no variation in the invasion growth rates among subpopulations of individuals. However, phenotypic, spatial, and temporal variation in vital rates can impact competitive outcomes [Levin, 1974, Warner and Chesson, 1985, Chesson, 1994, 2000b, Schreiber et al., 2011, Vasseur et al., 2011, Hart et al., 2016, Stump et al., 2021]. Variation in vital rates often results in variation in invasion growth rates among subpopulations of individuals within generations (spatial or phenotypic variation) or between generations (temporal variation). In these situations, the distribution of these invasion growth rates, not only the mean, can play a role in coexistence times. Understanding their role and how coexistence times depend on the nature of the variation (phenotypic vs. spatial vs. temporal) is a major challenge for future work. As our methodology for computing intrinsic coexistence is applicable to models accounting for this variation (see details in Appendix S8), it may provide a unified approach to tackling this challenge.

Some progress has been made on understanding the role of temporal variation on coexistence times [Pande et al., 2020b, Ellner et al., 2020, Pande et al., 2020a]. For example, using the methods presented here, [Ellner et al., 2020] found that the mean invasion growth rates for the lottery model [Warner and Chesson, 1985] are strongly correlated with intrinsic coexistence times. However, [Pande et al., 2020b,a] showed that ignoring the temporal variation in invasion growth rates can result in incorrect inferences about coexistence times. For example, greater environmental variation can increase mean invasion growth rates via the storage effect [Chesson, 1994], but can also lead to shorter coexistence times via the storage effect. Similar conclusions may also apply to within generational sources of variation. For example, increasing within-generational variation in fecundity decreases the persistence times of single species [Melbourne and Hastings, 2008]. We anticipate similar effects on coexistence times in our models.

The complexities we have identified in our study highlight the need to understand deterministic coexistence from both equilibrium-based and invasion-based approaches. Recent relevant equilibrium-based theories include Saavedra et al. [2017]‘s structural stability of feasible equilibria and Barabás et al. [2014]’s sensitivity analysis of stable equilibria. Both of these approaches provide insights into the range of demographic parameter values for which the equilibrium abundances remain positive i.e. feasible. Moreover, Saavedra et al. [2017] have developed feasibility metrics analogous to stabilizing niche differences and fitness differences. These developments raise the promising possibility of developing more informative, integrative metrics for species coexistence times.

## Acknowledgements

The authors thank the anonymous reviewers for their valuable feedback on earlier drafts of this manuscript. S.J.S. and J.M.L. were funded by the U.S. National Science Foundation Grants DMS-1716803 and DEB-2022213, respectively. N.J.B.K. was funded by U.S. National Science Foundation Grants DEB 1644641 and 2022810. O.G. acknowledges financial support provided by the Spanish Ministry of Economy and Competitiveness (MINECO) and by the European Social Fund through the Ramón y Cajal Program (RYC-2017-23666).

## Appendix S1 Model parameterization

In this Appendix, we derive the reduction of the annual plant with a seed bank in equation (3) to our deterministic model form in equation (1). Given that including the seed bank greatly complicates our analysis, yet ignoring it would unfairly bias competition in the system, we assume that seeds that ultimately germinate do so in the year after they are produced. If *s_i_* is the probability of a seed surviving and *g_i_* is its yearly germination probability, then the probability of the seed germinating in the next year is *g_i_*, two years is *g_i_*(1 – *g_i_*)*s_i_*, three years is 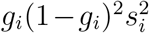. etc. Hence, the probability of a seed ultimately germinating 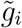 in (3) equals

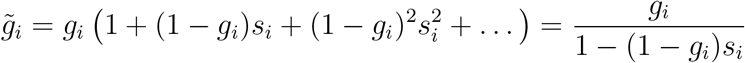

where the simplification follows from the geometric series equation 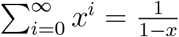 whenever |*x*| < 1.

Thus, the seed bank model in equation (3) reduces to:

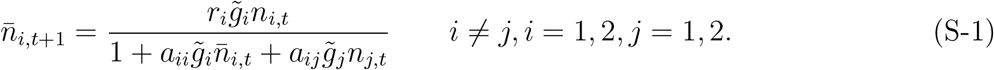

This equation is equivalent to model (1) after setting 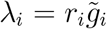 and 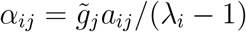. Therefore the coexistence times, invasion growth rates, niche overlap, and fitness differences can be calculated for the empirical system following the expressions outlined in the methods. Of the all possible 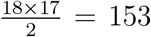 species pairs from 18 distinct species, only 90 species pairs had parameter estimates for which *r_i_* > 1 (i.e. the species could persist on its own), *α_ii_*, *α_ij_* are estimated and positive, and niche overlap *ρ* was less than one. Our analyses were performed on these 90 species pairs.

## Appendix S2 Deterministic approximation and extinction

In this Appendix, we prove two results. First, we prove that for any given time frame [0,*T*], the probability of stochastic model *n_t_* = (*n*_1,*t*_, *n*_2,*t*_) deviating from the mean field model 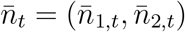 is arbitrarily small for 0 ≤ *t* ≤ *T* provided the habitat size *S* is sufficiently large. Second, we prove that, despite this strong correspondence over finite time frames, the stochastic models goes to extinction in finite time with probability one. To facilitate the presentation of these conclusions, define *f*(*n*) = (*f*_1_(*n*), *f*_2_(*n*)) where *f_i_*(*n*) = *n_i_*λ_*i*_(1 + *α_ii_n_i_* + *α_ij_n_j_*) where *i, j* ∈ {1,2} and *i* ≠ *j*. Let *M* = max{λ_1_/*α*_11_, λ_2_/*α*_22_}. Notice that *f_i_*(*n*) ≤ *M* for all *n* = (*n*_1_, *n*_2_).

For the first assertion, let *T* ≥ 1 and ε > 0 be given. Assume that 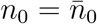. We will show that

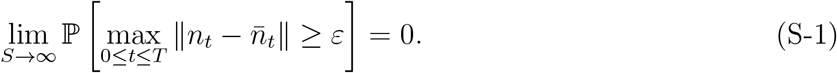

Continuity of *f* implies that there exists *δ* > 0 such that 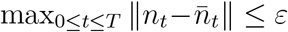 whenever ||*f*(*n*_*t*−1_) – *n_t_*|| ≤ *δ* for 1 ≤ *t* ≤ *T*. As the mean and variance of a Poisson random variable are equal, Chebyshev’s inequality implies

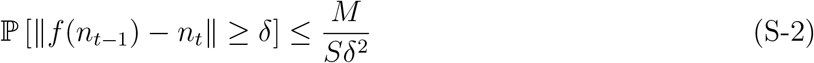

from which (S-1) follows.

For the second assertion, let

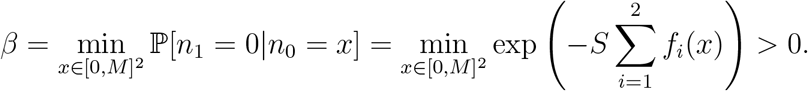

Next we use the following standard result in Markov chain theory [Durrett, 1996, Theorem 2.3 in Chapter 5].

### Proposition.

Let *X* be a Markov chain and suppose that

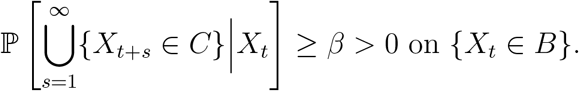

Then

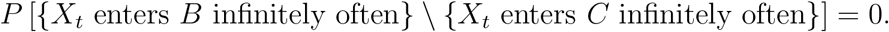

The proposition with *B* = [0, *M*]^2^ and *C* = {(0, 0)} implies that 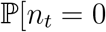 for some *t* ≥ 1] = 1.

## Appendix S3 Quasi-stationary distributions and coexistence times for large habitat sizes

In this Appendix, we prove the existence of quasi-stationary distributions for our individual-based model (2) and mathematically characterize the behavior of the intrinsic coexistence times times for larger habitat sizes *S*.

As our individual-based model (2) is an example of a nonlinear Poisson branching process (see, section 5.2 of Faure and Schreiber [2014] for a definition) whose mean-field dynamic (1) takes [0, ∞)^2^ into the compact set [0, λ_1_/(*α*_11_(λ_1_ – 1) × [0, λ_2_/(α_22_(λ_2_ – 1)) in one time step, Proposition 6.1 from Faure and Schreiber [2014] implies there is a unique quasi-stationary distribution of (2).

Now, assume that the deterministic model (1) predicts coexistence i.e. 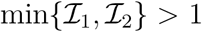. Then, Cushing et al. [2004] proved that the coexistence equilibrium 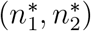 of the deterministic model is globally stable i.e. all initial conditions of the deterministic mean-field model supporting both species asymptotically approach this equilibrium. Consequently, there is only one invariant probability measure supported in (0, ∞)^2^ for the deterministic model–the Dirac measure at this coexistence equilibrium. Lemma 3.9 and Theorem 5.4 of Faure and Schreiber [2014] imply that (i) the quasi-stationary distributions converge in the weak* topology to this Dirac measure as S ↟ ∞ and (ii) there exist constants *α* > 0, ✶ > 0 such that the intrinsic coexistence time is ≥ *α* exp(✶*S*) for sufficiently large habitat sizes *S*.

Now, assume that the deterministic model (1) predicts competitive exclusion, say species 1 asymptotically excludes species 2. Let *π^S^*(*N*_1_, *N*_2_) be the QSD for the habitat size *S* and let 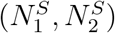 be random variables whose distribution is given by *π^S^*. Then the arguments used to prove Theorem 2.7 in Faure and Schreiber [2014] can be modified to show that the quasi-stationary distributions converge in the weak* topology to the Dirac measure at the single species equilibrium (1/*α*_11_, 0) as *S* → ∞. In particular, given any *ε* > 0, there exists a habitat size *S*(*ε*) > 0 such that

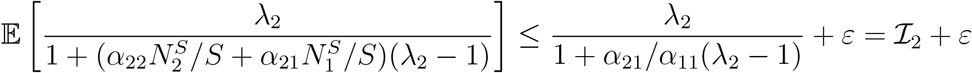

for *S* ≥ *S*(*ε*). It follows from the definition of quasi-stationarity that

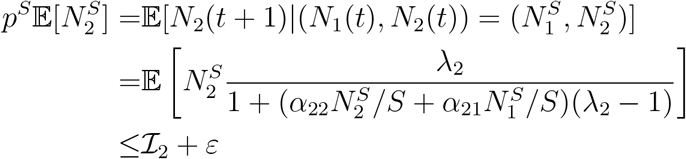

for *S* ≥ *S*(*ε*). Hence, the intrinsic coexistence time is less than

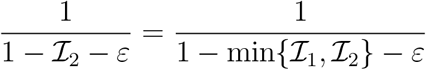

for *S* ≥ *S*(*ε*) and in the limit of *S* → ∞ we get that the intrinsic coexistence time is less than 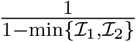

## Appendix S4 Global sensitivity analysis of coexistence times

**Figure S1.**
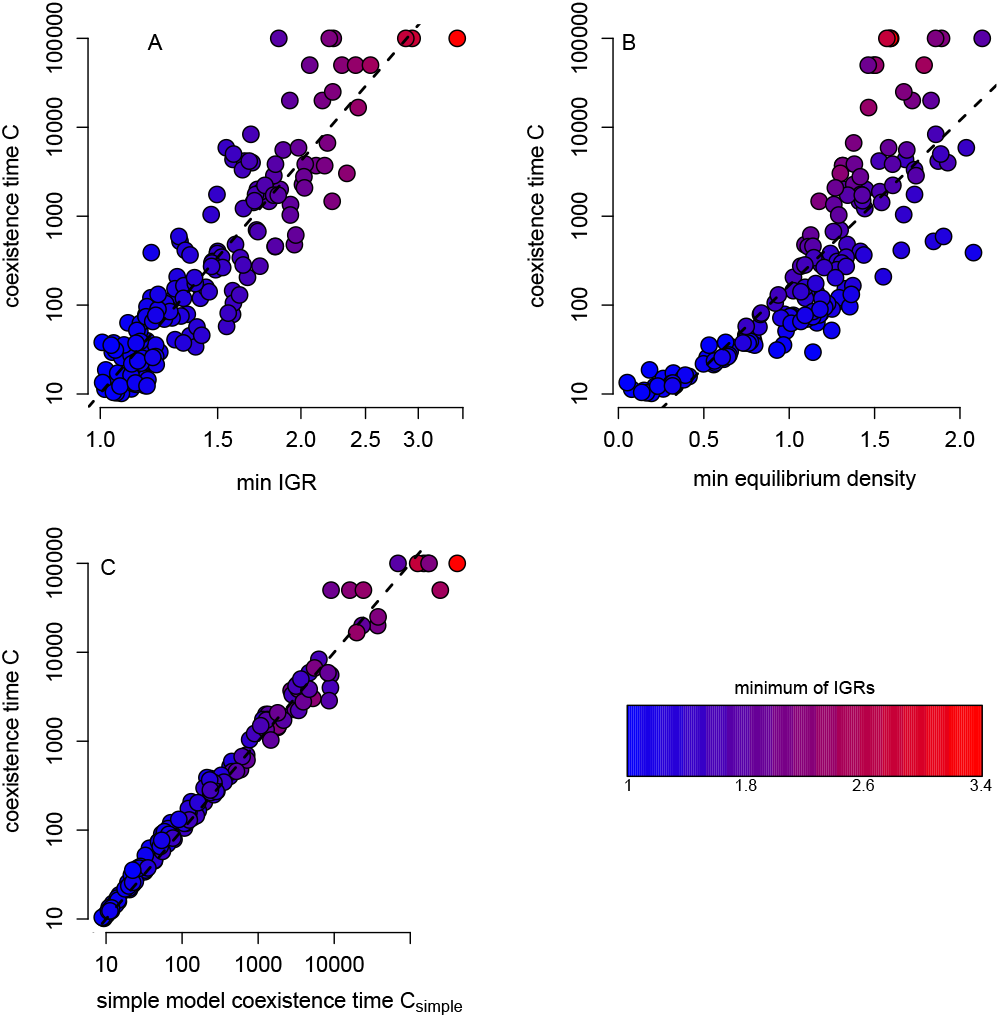
200 random parameter samples with λ_*i*_ uniformly distributed in [1.1, 2], α_*ii*_ uniformly distributed in [0.2, 0.5], *α_ij_* with *j* = *i* uniformly distributed in [0, *α_ii_*]; the last restriction ensures deterministic coexistence i.e. 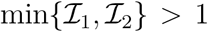. In panel A, a linear regression of log_10_*C* against 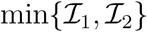 explained approximately 82% of the variance (adjusted-R^2^ of 0.824) with estimated intercept 1.00761 and slope 8.68233. In panel B, a linear regression of log_10_*C* against 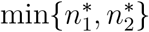 explained approximately 70% of the variance (adjusted-R^2^ of 0.717) with estimated intercept 0.36795 and slope 1.85465. In panel C, a linear regression of log_10_*C* against log_10_*C*_simple_ with intercept 0 explained nearly all of the variance (adjusted-R^2^ of 0.9963) with estimated slope 1.006993.

## Appendix S5 Analysis of equilibrium densities

This appendix analyzes how the equilibria densities of the deterministic model change with niche overlap*ρ* and the fitness ratio *k_j_*/*k_j_*. Recall that at the coexistence equilibrium, the density of species *i* is given by

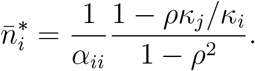

This expression is decreasing linearly with the fitness ratio *k_j_*/*k_j_* associated with species *j* i.e. as species *j* becomes a better competitor, the equilibrium density of species *i* decreases. The derivative of 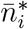 with respect to *ρ* equals

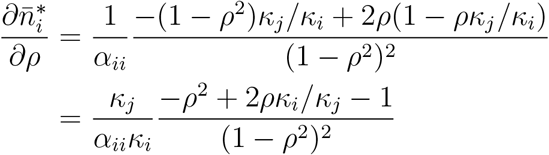

When *k_i_*/*j_j_* < 1 (i.e. species *i* is the fitness inferior), the numerator is negative for all *ρ* ≥ 0. Hence, the equilibrium density of the fitness inferior decreases with niche overlap, as claimed in the main text. In contrast, when *κ_i_*/*κ_j_* > 1 (i.e. species *i* is the fitness superior), the numerator of this derivative is negative for 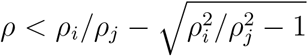 and positive for 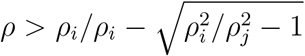. In particular, as coexistence only occurs, in this case, for *ρ* < *ρ_j_*/*κ_i_κ_i_* and

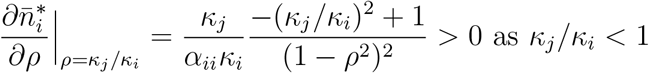

the equilibrium density of the fitness superior decreases and then increase with niche overlap.

## Appendix S6 Effects of *ρ* and *κ_j_*/*κ_j_* on coexistence times

In this Appendix, we explored the effects of *ρ* and *κ_i_*/*κ_j_* of the fitness superior for 5 of deterministically coexisting pairs; the results for the other two species pairs (#1,#4) are shown in Figs. 4,5 in the main text. Red arrows correspond to base empirical value.

**Figure S1.**
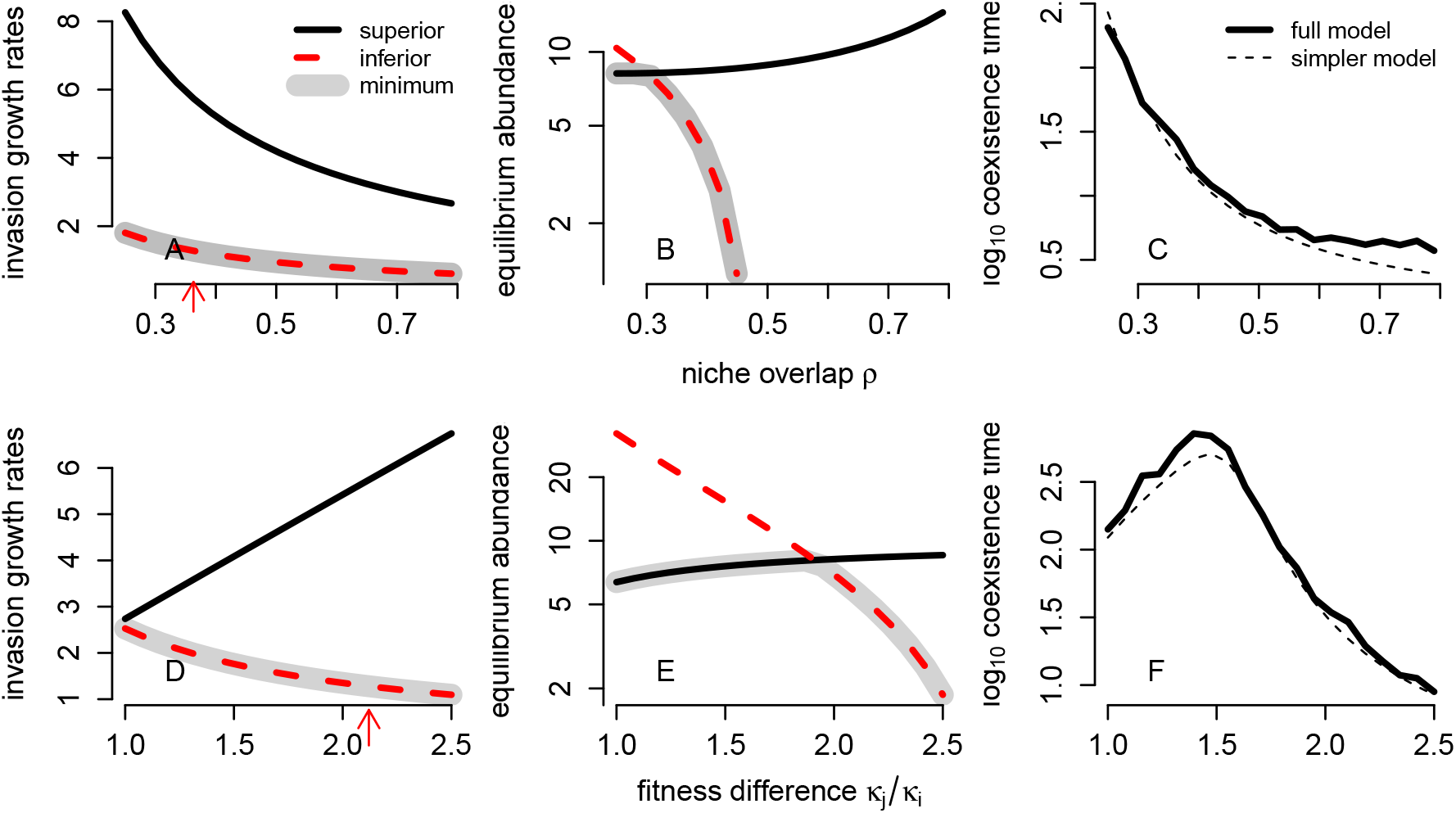
Species Pair #2.

**Figure S2.**
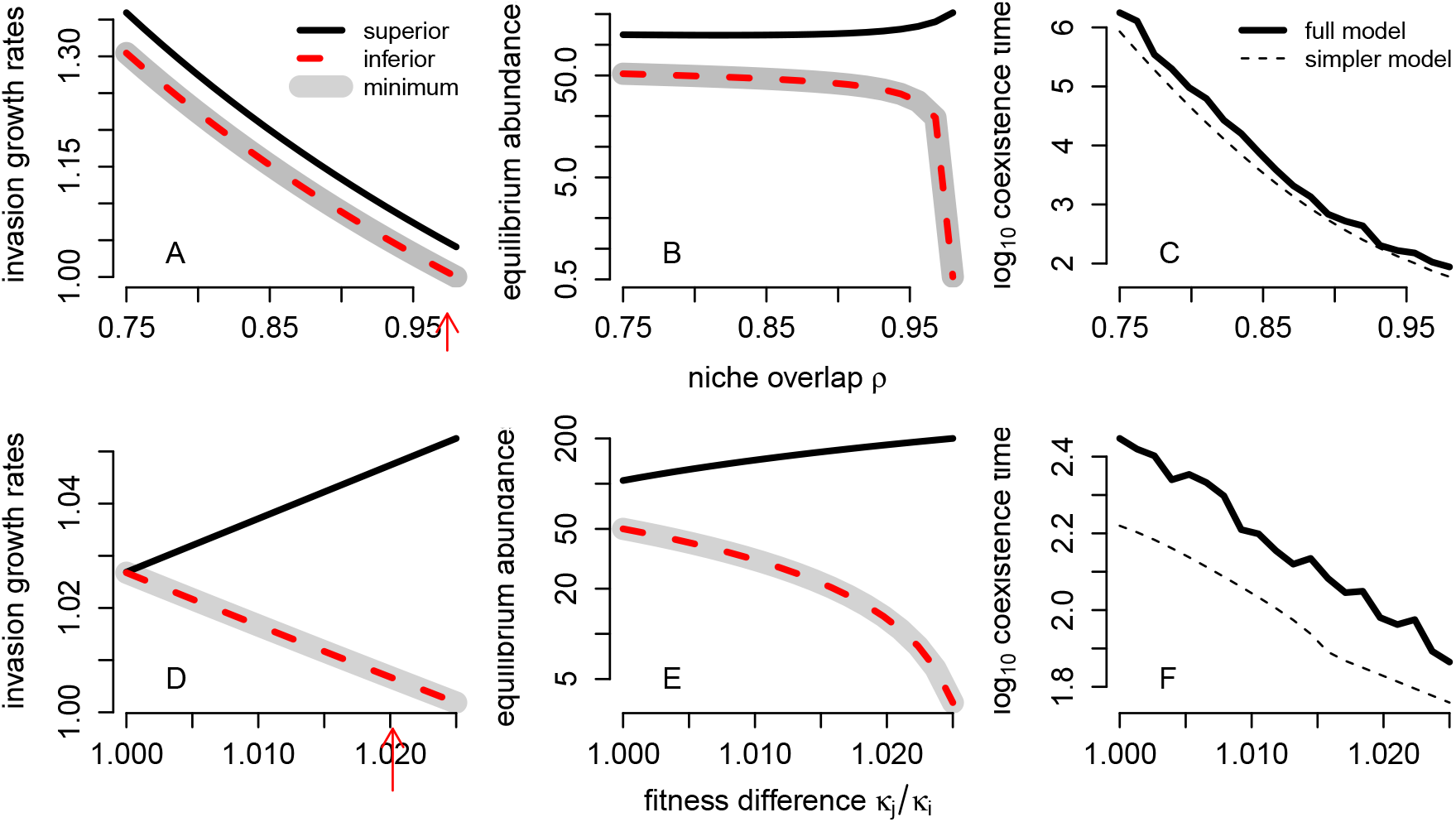
Species Pair #3.

**Figure S3.**
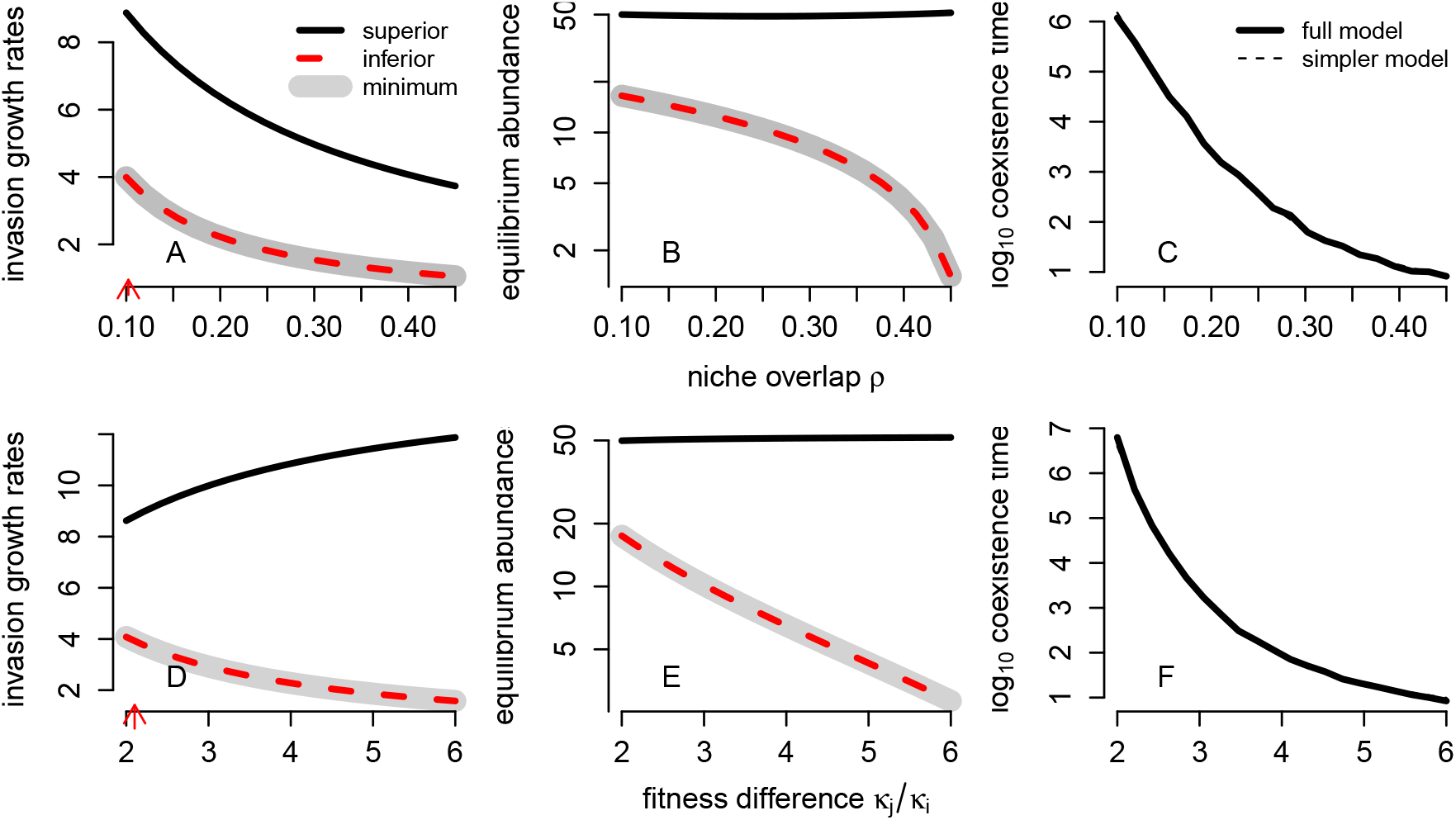
Species Pair #5.

**Figure S4.**
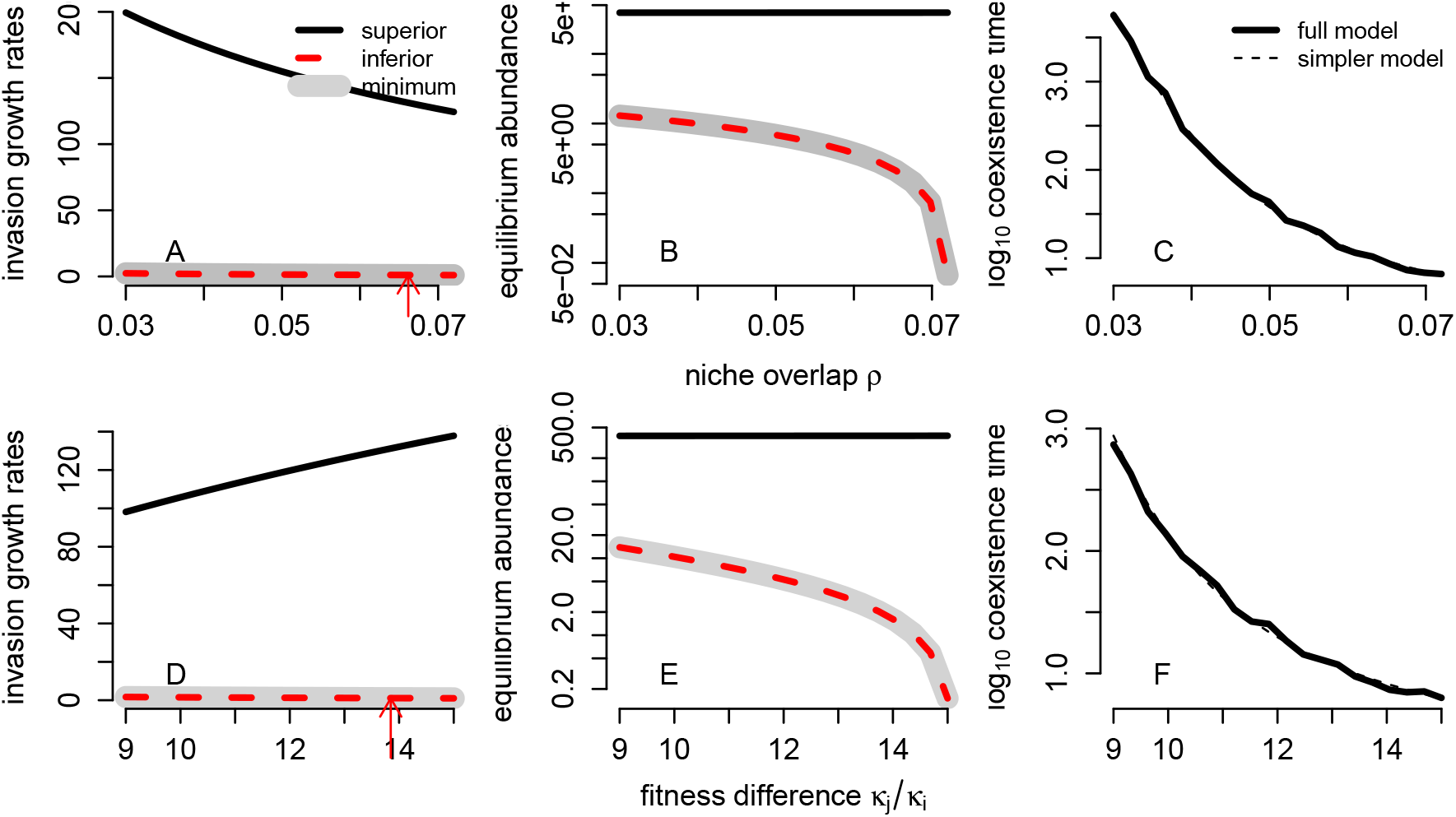
Species Pair #6.

**Figure S5.**
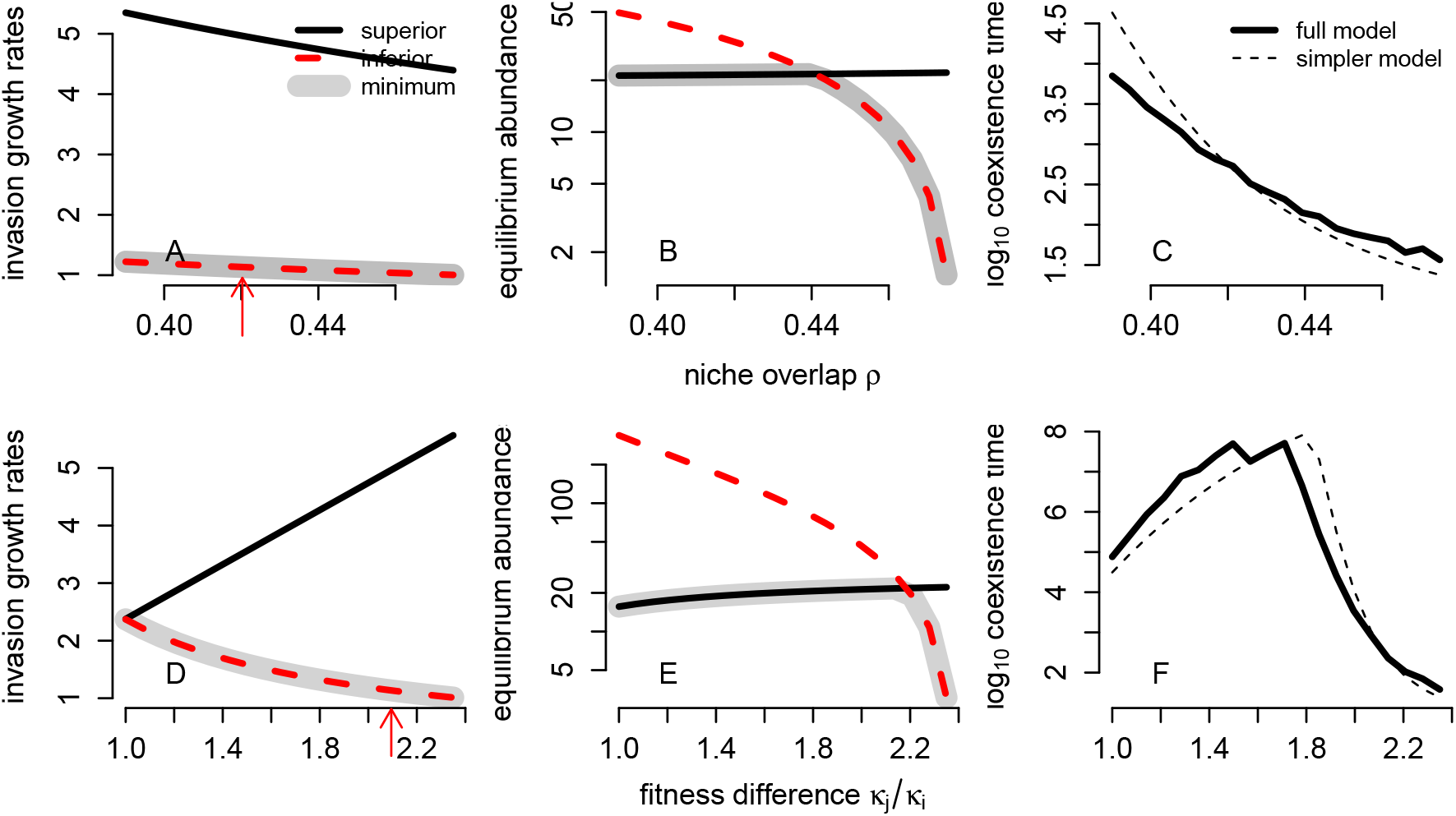
Species Pair #8

## Appendix S7 Coexistence times for the simplified model

To determine the coexistence times for the simplified model, let *P_i_* be the mean persistence time for each species. As these times are geometrically distributed, the probability of extinction in each time step is 1/*P_i_* for species. As the two species dynamics are independent in the simplified model, the probability that neither goes extinct in a time step is (1 – 1/*P*_1_)(1 – 1/*P*_2_), and the probability at least one goes extinct in a time step is 1 – (1 – 1/*P*_1_)(1 – 1/*P*_2_). Hence, the mean time to losing one of the species is

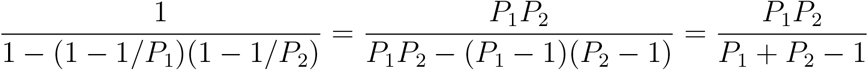

as claimed in the main text. For *P*_1_ » 1 and *P*_2_ » 1, we have

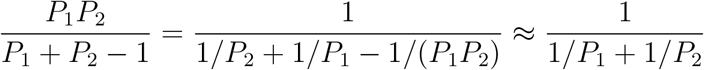

which equals 1/2 of the harmonic mean of the 1/*P_i_* as claimed in the main text.

Using these expressions, we can derive various insights about the two species model. First, we will show that 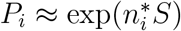 whenever 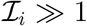. Indeed, for *N_i,t_*, ≥ 1 in (8)

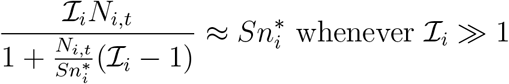

Thus, whenever *N_i,t_*, ≥ 1,

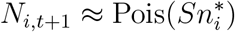

and the probability of extinction is approximately 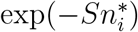. As these approximations hold whenever *N_i,t_*, ≥ 1, they describe both the quasi-stationary distribution and quasi-extinction probability. Hence, *P_i_* ≈ exp(–*n_i_S*).

Next, suppose that both species have 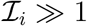 (equivalently 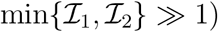. Then

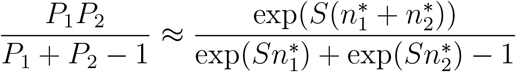

Subject to the constraint that 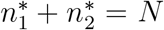 (i.e. a maximal community size), one can show using Lagrange multipliers that the coexistence time is maximized when 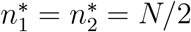.

Finally, suppose that 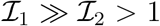. Then 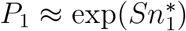 and 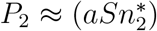 for some 0 < *a* < 1. Subject to the constraint that 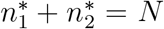 (i.e. a maximal community size), one can show using Lagrange multipliers that the coexistence time is maximized when 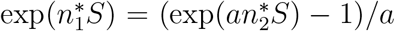 subject to the constraint 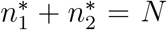. As (exp(*ax*) – 1)/*a* > exp(*x*) – 1 for all *x* > 0, it follows that one must have that 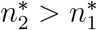.

## Appendix S8 Quasi-stationary distributions for spatially structured communities in fluctuating environments

This Appendix describes how the methods used in the main text extend to ecological models accounting for any number of species, any types of species interactions, discrete spatial structure, and environmental fluctuations. After describing the model with and without demographic stochasticity, we present conditions that ensure the existence and uniqueness of a quasi-stationary distribution. Then we describe how the algorithm of Aldous et al. [1988] can be used to numerically estimate the quasi-stationary distributions. We conclude by discussing how the scaling of coexistence times with the community size *S* depend on the nature of the temporal fluctuations. While we focus on a model using Poisson updates for clarity, the same ideas apply to model with mixtures of multinomials and negative binomials.

### Spatially structured community dynamics in fluctuating environments

The mean field model allows for *k* species and *ℓ* spatial patches. For simplicity, assume that all species have synchronized generations (e.g. competing annual plants, host-parasitoid interactions) and a time step correspond this generation time. Let *n_ij_* denote the density of species *i* in patch *j* and 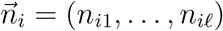 the vector of species *i*’s densities across the patches. Let 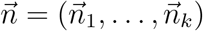 be the state of the community. To model the environmental fluctuations, let *ξ*_1_, *ξ*_2_,… be an ergodic, stationary sequence of random vectors that represent the fluctuations in the species parameters. For each species *i*, let 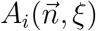 be a non-negative, *ℓ* × *ℓ* projection matrix such that 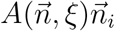 is species *i*’s state in the next time step when the current community state is 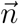 and the current environmental state is *ξ*. The entries of 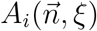 correspond to reproduction and dispersal that may depend on the state of the community and the environment. The mean field model is

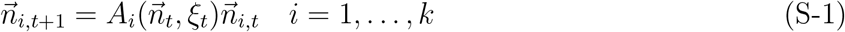

To account for demographic stochasticity, we assume that births are Poisson distributed and dispersal occurs via multinomial sampling. As discussed in Faure and Schreiber [2014, Section 5.2], these assumptions imply that updates of each species in each patch are given by independent Poisson random variables whose means are given by mean field model. Namely, if *S* is the community size and 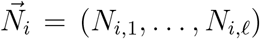 are the abundances of species *i* across the patches, then the stochastic model is

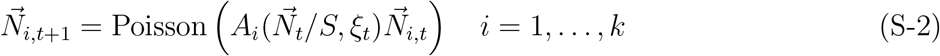

A standard argument discussed in Schreiber [2017] implies that *N_t_* goes to zero (i.e. all species go extinct) in finite time with probability one. In particular, the process enters the extinction set

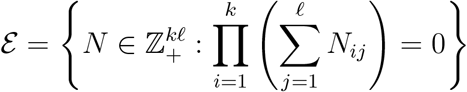

in finite time with probability one where 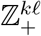 refers to the non-negative integer lattice of dimension *kℓ* i.e. abundances of *k* species in *ℓ* patches.

### Quasi-stationary distributions and their estimation

Let 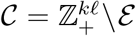 be the set of coexistence states. For a given community size *S*, a quasi-stationary distribution for (S-2) is a distribution 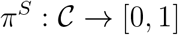 on the non-extinction states (i.e. 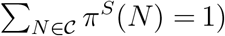 if there exists λ^*S*^ ∈ (0,1) such that 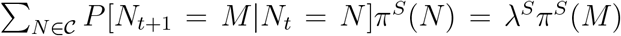. λ^*S*^ corresponds to the probability of all species coexisting over one generation given the model is in the quasi-stationary state. Hence, 1/(1 – λ^*S*^) is the intrinsic coexistence time. The argument in the proof of Faure and Schreiber [2014, Proposition 6.3] can be used to show that there exists a unique quasi-stationary distribution whenever the mean field model is a compact map (i.e. there exists a compact set *K* such that *n*_*t*+1_ ∈ *K* for all *n_t_* in (S-1)) and *ξ* takes values in a compact set.

To numerically estimate the quasi-stationary distribution using the Aldous et al. [1988] algorithm, assume that the stationary sequence *ξ*_1_, *ξ*_2_,… is given by a Markov process with transition operator *T* i.e. *T*(*ξ,B*) = *P*(*ξ*_*t*+1_ ∈ *B*|*ξ*_*t*_ = *ξ*) for any measurable set *B* of environmental states. To estimate *π^S^*, we simulate a modified version of the process *ξ_t_, N_t_*. Namely, given *ξ_t_, N_t_* find *ξ*_*t*+1_, *N*_*t*+1_ by two steps

1. Draw a multivariate Poisson *N*_*t*+1_ as given by (S-2) and draw *ξ*_*t*+1_ independently from the law of the probability measure *T*(*ξ*_*t*_, *dξ*).
2. If 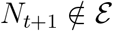, then use (*N*_*t*+1_, *ξ*_*t*+1_). If 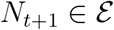, then replace (*ξ*_*t*+1_, *N*_*t*+1_) with a uniform random draw from the past states (*ξ*_0_, *N*_0_), (*ξ*_1_, *N*_1_)…, (*ξ_t_*, *N_t_*) i.e. with probability 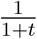 replace with state (*ξ_s_*, *N_s_*) for 0 ≤ *s* ≤ *t*.

For a sufficiently long run of this modified process, the empirical distribution of *N_t_* approximates the quasi-stationary distribution i.e. the fraction of time spent the modified process *N_t_* spends in community state *N* approximates *π^S^*(*N*). Furthermore, the faction of time step 2 is taken approximates the extinction probability 1 – λ^*S*^.

### Scaling of coexistence times with community size *S*

When the environmental dynamic is periodic (i.e. *ξ*_1_, *ξ*_2_,… is periodic) and the mean field model (S-1) has a positive attractor, the argument in the proof of Faure and Schreiber [2014, Theorem 2.7] implies that the coexistence time 1/(1 – λ^S^) increases exponentially with the community size S. For stochastic environments (e.g. *ξ*_1_, *ξ*_2_,… are i.i.d.), the scaling of the coexistence time with community size *S* is more subtle [Guillin et al., 2019, Ellner et al., 2020]. Theoretical and simulation results suggest that the coexistence time scale (at least) polynomially with *S* whenever the mean field model is stochastically persistent but can be exponential with the positive stationary distribution is bounded away from extinction.

## References

P. Adler, J. HilleRisLambers, and J. Levine. A niche for neutrality. Ecology Letters, 10:95–104, 2007.

P. Adler, S. Ellner, and J. Levine. Coexistence of perennial plants: an embarrassment of niches. Ecology Letters, 13:1019–1029, 2010.

P. B. Adler and J. M. Drake. Environmental variation, stochastic extinction and competitive coexistence. The American Naturalist, 172, 2008.

D. Aldous, B. Flannery, and J. Palacios. Two applications of urn processes the fringe analysis of search trees and the simulation of quasi-stationary distributions of Markov chains. Probability in the Engineering and Informational Sciences, 2:293–307, 1988.

G. Barabás, L. Pásztor, G. Meszéna, and A. Ostling. Sensitivity analysis of coexistence in ecological communities: theory and application. Ecology Letters, 17(12):1479–1494, sep 2014.

M. Benaïm and B. Cloez. A stochastic approximation approach to quasi-stationary distributions on finite spaces. Electronic Communications in Probability, 20, 2015.

R. J. H. Beverton and S. J. Holt. On the dynamics of exploited fish populations, volume 2(19) of *Fisheries Investigation Series*. Ministry of Agriculture, Fisheries and Food, London, UK, 1957.

M. Boyce. Population viability analysis. Annual Review of Ecology & Systematics, 23:481–497, 1992.

I. Carroll, B. Cardinale, and R. Nisbet. Niche and fitness differences relate the maintenance of diversity to ecosystem function. Ecology, 92:1157–1165, 2011.

P. Chesson. Macarthur’s consumer-resource model. Theoretical Population Biology, 37:26–38, 1990.

P. Chesson. Multispecies competition in variable environments. Theoretical Population Biology, 45: 227–276, 1994.

P. Chesson. Mechanisms of maintenance of species diversity. Annual Review of Ecology and Systematics, 31:343–366, 2000a.

P. Chesson. General theory of competitive coexistence in spatially-varying environments. Theoretical Population Biology, 58:211–237, 2000b.

P. Chesson. *Ecological Systems*, chapter Species Competition and Predation, pages 223–256. Springer New York, New York, NY, 2013.

P. Chesson. Updates on mechanisms of maintenance of species diversity. Journal of Ecology, 106: 1773–1794, 2018.

C. Chu and P. Adler. Large niche differences emerge at the recruitment stage to stabilize grassland coexistence. Ecological Monographs, 85:373–392, 2015.

J. Cushing, S. Levarge, N. Chitnis, and S. Henson. Some discrete competition models and the competitive exclusion principle†. Journal of Difference Equations and Applications, 10:1139–1151, 2004.

R. Durrett. Probability: Theory and examples. Duxbury Press, Belmont, CA, 1996.

S. Ellner, R. Snyder, and P. Adler. How to quantify the temporal storage effect using simulations instead of math. Ecology Letters, 19:1333–1342, 2016.

S. Ellner, R. Snyder, P. Adler, G. Hooker, and S. Schreiber. Technical Comment on Pande et al. (2020): Why invasion analysis is important for understanding coexistence. Ecology Letters, 23 (11):1721–1724, 2020.

S. P. Ellner, R. E. Snyder, P. B. Adler, and G. Hooker. An expanded modern coexistence theory for empirical applications. Ecology Letters, 22:3–18, 2019.

W. J. Ewens. Mathematical Population Genetics I. Theoretical Introduction, volume 27. Springer Science & Business Media, 2012.

M. Faure and S. J. Schreiber. Quasi-stationary distributions for randomly perturbed dynamical systems. Annals of Applied Probability, 24:553–598, 2014.

A. Gabel, B. Meerson, and S. Redner. Survival of the scarcer. Phys Rev E Stat Nonlin Soft Matter Phys, 87:010101, 2013.

B. Gilbert and J. Levine. Plant invasions and extinction debts. Proceedings of the National Academy of Sciences, 110:1744–1749, 2013.

O. Godoy and J. M. Levine. Phenology effects on invasion success: insights from coupling field experiments to coexistence theory. Ecology, 95:726–736, 2014.

O. Godoy, N. Kraft, and J. Levine. Phylogenetic relatedness and the determinants of competitive outcomes. Ecology Letters, 17:836–844, 2014.

A. Gómez-Corral and M. López García. Extinction times and size of the surviving species in a two-species competition process. Journal of Mathematical Biology, 64:255–289, 2012.

T. N. Grainger, J. M. Levine, and B. Gilbert. The invasion criterion: a common currency for ecological research. Trends in Ecology and Evolution, 2019.

V. Grimm and C. Wissel. The intrinsic mean time to extinction: a unifying approach to analysing persistence and viability of populations. Oikos, 105:501–511, 2004.

A. Guillin, A. Personne, and E. Strickler. Persistence in the moran model with random switching. arXiv preprint arXiv:1911.01108, 2019.

S. Hart, S. Schreiber, and J. Levine. How variation between individuals affects species coexistence. Ecology Letters, 19:825–838, 2016.

S. P. Hart, R. P. Freckleton, and J. M. Levine. How to quantify competitive ability. Journal of Ecology, 106:1902–1909, 2018.

J. Hofbauer and K. Sigmund. Evolutionary games and population dynamics. Cambridge University Press, 1998.

S. P. Hubbell. The unified neutral theory of biodiversity and biogeography, volume 32 of Monographs in population biology. Princeton University Press, Princeton, New Jersey, 2001.

S. P. Hubbell and R. B. Foster. Biology, chance and history and the structure of tropical rainforest tree communities, pages 314–329. Harper and Row, New York, 1986.

G. Hutchinson. The paradox of the plankton. The American Naturalist, 95:137–145, 1961.

P. Jagers. A plea for stochastic population dynamics. Journal of Mathematical Biology, 60:761–764, 2010.

A. M. Kramer and J. M. Drake. Time to competitive exclusion. Ecosphere, 5:art52, 2014.

R. Lande, S. Engen, and B.-E. Saether. *Stochastic population dynamics in ecology and conservation*. Oxford series in ecology and evolution. Oxford University Press, 2003.

P. H. Leslie and J. C. Gower. The properties of a stochastic model for two competing species. Biometrika, 45:316–330, 1958.

A. Letten, P. Ke, and T. Fukami. Linking modern coexistence theory and contemporary niche theory. Ecological Monographs, 87:161–177, 2017.

S. Levin. Dispersion and population interactions. The American Naturalist, 108:207–228, 1974.

J. M. Levine and J. Hille Ris Lambers. The importance of niches for the maintenance of species diversity. Nature, 461:254–257, 2009.

S. Li, J. Tan, X. Yang, C. Ma, and L. Jiang. Niche and fitness differences determine invasion success and impact in laboratory bacterial communities. The ISME Journal, 13:402–412, 2019.

R. H. Macarthur and R. Levins. The limiting similarity, convergence, and divergence of coexisting species. American Naturalist, 101:377–385, 1967.

R. M. May. Stability and Complexity in Model Ecosystems, 2nd edn. Princeton University Press, Princeton, 1975.

C. Meine. It’s about time: Conservation biology and history. Conservation Biology, 13:1–3, 2 1999.

B. Melbourne and A. Hastings. Extinction risk depends strongly on factors contributing to stochas-ticity. Nature, 454:100, 2008.

S. Méléard and D. Villemonais. Quasi-stationary distributions and population processes. Probability Surveys, 9:340–410, 2012.

E. Mordecai, K. Gross, and C. Mitchell. Within-host niche differences and fitness trade-offs promote coexistence of plant viruses. The American Naturalist, 187(1):E13–E26, 2016.

H. Muller-Landau and M. Visser. How do lianas and vines influence competitive differences and niche differences among tree species? concepts and a case study in a tropical forest. Journal of Ecology, 107:1469–1481, 2019.

A. Narwani, M. A. Alexandrou, T. H. Oakley, I. T. Carroll, and B. J. Cardinale. Experimental evidence that evolutionary relatedness does not affect the ecological mechanisms of coexistence in freshwater green algae. Ecology Letters, 16:1373–1381, 2013.

O. Ovaskainen and B. Meerson. Stochastic models of population extinction. Trends in Ecology and Evolution, 25:643–652, 2010.

J. Pande, T. Fung, R. Chisholm, and N. Shnerb. Invasion growth rate and its relevance to persistence: a response to Technical Comment by Ellner et al. Ecology Letters, 23:1725–1726, 2020a.

J. Pande, T. Fung, R. Chisholm, and N. M. Shnerb. Mean growth rate when rare is not a reliable metric for persistence of species. Ecology Letters, 23:274–282, 2020b.

G. Reuter. Competition processes, pages 421–430. University of California Press, Berkeley, 1961.

S. Saavedra, R. P. Rohr, J. Bascompte, O. Godoy, N. J. B. Kraft, and J. M. Levine. A structural approach for understanding multispecies coexistence. Ecological Monographs, 87(3):470–486, 6 2017.

S. Schreiber, M. Benaïm, and K. A. S. Atchadé. Persistence in fluctuating environments. Journal of Mathematical Biology, 62:655–683, 2011.

S. J. Schreiber. Criteria for *C^r^* robust permanence. Journal of Differential Equations, 162:400–426, 2000.

S. J. Schreiber. Coexistence in the face of uncertainty. In Recent Progress and Modern Challenges in Applied Mathematics, Modeling and Computational Science, pages 349–384. Springer, 2017.

S. J. Schreiber, M. Yamamichi, and S. Y. Strauss. When rarity has costs: coexistence under positive frequency-dependence and environmental stochasticity. Ecology, 100:e02664, 2019.

L. Shoemaker, L. Sullivan, I. Donohue, J. Cabral, R. Williams, M. Mayfield, J. Chase, C. Chu, W. Harpole, A. Huth, et al. Integrating the underlying structure of stochasticity into community ecology. Ecology, 101:e02922, 2020.

C. Song, G. Barabás, and S. Saavedra. On the consequences of the interdependence of stabilizing and equalizing mechanisms. The American Naturalist, 194(5):627–639, 2019.

J. Spaak and F. De Laender. Intuitive and broadly applicable definitions of niche and fitness differences. Ecology Letters, 23:1117–1128, 2020.

S. Stump, C. Song, S. Saavedra, J. Levine, and D. Vasseur. Synthesizing the effects of individuallevel variation on coexistence. Ecological Monographs, page e1493, 2021.

D. Vasseur, P. Amarasekare, V. Rudolf, and J. Levine. Eco-evolutionary dynamics enable coexistence via neighbor-dependent selection. The American Naturalist, 178:E96–E109, 2011.

M. Vellend. Conceptual synthesis in community ecology. Quarterly Review of Biology, 85:183–206, 2010.

C. Wainwright, J. HilleRisLambers, H. Lai, X. Loy, and M. Mayfield. Distinct responses of niche and fitness differences to water availability underlie variable coexistence outcomes in semi-arid annual plant communities. Journal of Ecology, 107:293–306, 2019.

R. Warner and P. Chesson. Coexistence mediated by recruitment fluctuations: A field guide to the storage effect. The American Naturalist, 125:769–787, 6 1985.

A. Watkinson. Density-dependence in single-species populations of plants. Journal of Theoretical Biology, 83(2):345 – 357, 1980.

V. Zepeda and C. Martorell. Fluctuation-independent niche differentiation and relative non-linearity drive coexistence in a species-rich grassland. Ecology, 100(8):e02726, 2019.

